# Amino-alcohol Bio-conjugate of Naproxen Exhibits Anti-inflammatory Activity through NF-κB Signaling Pathway

**DOI:** 10.1101/2020.01.10.901223

**Authors:** Mahua Rani Das, Anindyajit Banerjee, Sourav Sarkar, Joydeb Majumder, Saikat Chakrabarti, Siddhartha Sankar Jana

**Affiliations:** Department of Biological Chemistry; Indian Association for the Cultivation of Science; 2A & 2B, Raja Subodh Chandra Mallick Road; Jadavpur, Kolkata, West Bengal 700032, India; Department of Biology of Cancer & Other Diseases; National Centre for Cell Science; NCCS Complex; University of Pune Campus; University Road, Ganeshkhind, Pune, Maharashtra 411007, India; Structural Biology and Bio-informatics Division; CSIR-Indian Institute of Chemical Biology; 4, Raja Subodh Chandra Mallick Road; Poddar Nagar; Jadavpur, Kolkata, West Bengal 700032, India; Department of Bioinformatics, Tata Translational Cancer Research Centre, Tata Medical Center, Kolkata-700156, India; Director’s Research Unit, Indian Association for the Cultivation of Science; 2A & 2B, Raja Subodh Chandra Mallick Road; Jadavpur, Kolkata, West Bengal 700032, India; Department of Organic Chemistry, Indian Association for the Cultivation of Science; 2A & 2B, Raja Subodh Chandra Mallick Road; Jadavpur, Kolkata, West Bengal 700032, India; Department of Chemistry, Purdue University Centre for Drug Discovery, Purdue University, West Lafayette, IN 47906, USA

**Author notes:** **Corresponding Authors** I. Dr. Mahua Rani Das II. Dr. Saikat Chakrabarti.

**Keywords:** Naproxen sodium, Hydrogel, Naproxen Bio-Conjugates, Inflammation, Drug delivery

## Abstract

Naproxen sodium (Ns) is a well-known synthetic compound used vastly as a nonsteroidal anti-inflammatory drug (NSAID). Previously, we demonstrated that the hydrogels, made of Ns bio-conjugate (NBC), either with amino alcohol (NBC-1 and 2) or amino acid (NBC-3 and 4), displayed anti-inflammatory properties better than Ns, but the effectiveness was not well explored. Here, we investigated that NBC-2, conjugated with γ-amino alcohol, significantly reduces the expression of pro-inflammatory proteins; such as inducible nitric oxide synthase (iNOS) and cyclooxygenase isoform-2 (COX-2), and the production of pro-inflammatory mediator nitric oxide (NO) in lipopolysaccharide and interferon-γ (LPS/IFN-γ) treated mouse macrophage RAW 264.7 cells, compared with bio-conjugation of β-amino alcohol (NBC-1) or β-amino acid (NBC-3) or α-amino acid NBC (NBC-4). NBC-2 decreases the nuclear localization of transcription factors NF-κB by stabilizing its cytoplasmic functional inhibitor IκBα. Moreover, NBC-2 also displays selective inhibitory effect towards COX-1 enzyme activity as determined by enzymatic assay and *in silico* molecular docking analysis. Thus, we suggest that NBC-2 may be used as a self-delivery anti-inflammatory drug as compared with other NBCs.

**TOC Graphic:** 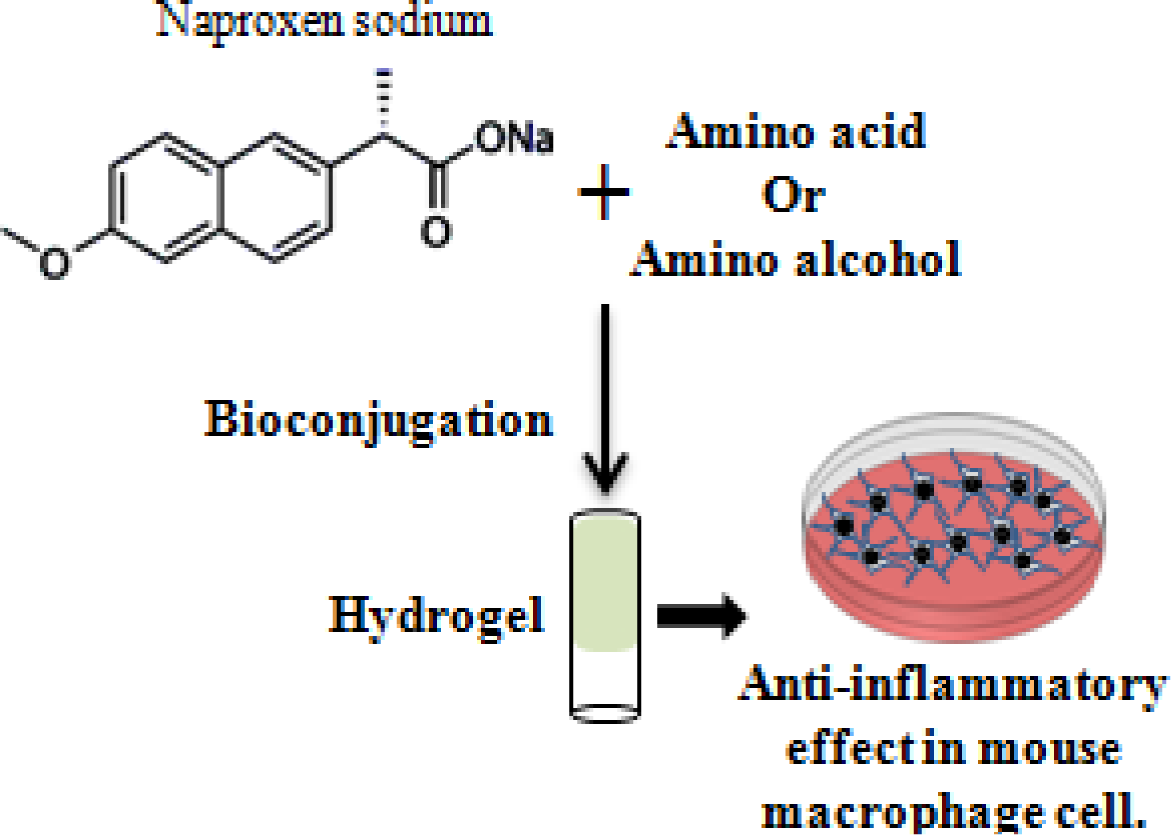

## Introduction

Nonsteroidal anti-inflammatory drugs (NSAIDs) are a class of drugs that provide analgesic and antipyretic effects at lower dose; whereas at higher dose, they show anti-inflammatory effects.^1^ Generally, NSAIDs act by inhibiting the pro-inflammatory protein cyclooxygenase (COX) activity. There are two isoforms of COX; one is expressed constitutively (COX-1), whereas the second is inducible (COX-2).^2^ Both isoforms convert arachidonic acid (AA) to prostaglandins (PGs) and a potent platelet aggregating agent thromboxane A4, resulting in inflammation by increased vascular permeability and vasodilation.^3^ When the body encounters any foreign stimulatory molecule, the macrophages play an important role in nonspecific defense (innate immunity) mechanisms and also help to initiate specific defense mechanisms (adaptive immunity) by releasing growth factors, cytokines, and lipid mediators such as prostaglandins and leukotrienes, which help to recruit of other immune cells, such as B-cells and T-cells, resulting in the regulation of immune response and inflammation.^4^ Naproxen (Np) is one of the oldest and largest selling non-selective NSAIDs that inhibit both COX isoforms with comparable IC_50_ value.^5^ But, according to Grosser et al., naproxen is a more potent inhibitor of COX-2 than COX-1.^6^ Most of the known NSAIDs show gastrointestinal bleeding and increased cardiovascular (CV) problems, but Np has the most favorable cardiovascular risk profile.^7^ On the other hand, Np inhibits colonic adenocarcinoma by suppressing the p65/β-catenin/cyclin D1 signaling pathway in rats.^8^ When administered in a combination therapy with calcitriol (the hormonally active form of vitamin D_3_), Np was found to be safe and effective in treating progressive prostate cancer.^9^ Naproxen sodium (Ns), sodium salt form of naproxen (Np), is more effective in the treatment of osteoarthritis of the knee as compared to other NSAIDs. It is also used to treat rheumatic diseases such as rheumatoid arthritis, osteoarthritis, ankylosing spondylitis and nonarticular rheumatism.^10^

Poor water solubility of Np leads low bioavailability.^11^ High dosages of Np may cause gastrointestinal and renal problems, which is overcome by converting Np (MW 230.0943) into hydrogelator-based self-delivery systems. Supramolecular hydrogels, derived from self-assembly of low molecular weight hydrogelators (LMWHs, MW < 3000) have recently drawn considerable attention for their wide range of applications in biological science, e.g. in tissue engineering,^12,13^ regenerative medicine,^14^ wound healing,^15^ drug delivery,^16–20^ biomineralization,^21^ inhibitors of cancer cells,^22^ 3D cell cultures etc.^23,24^ Therefore, in recent times significant efforts have been directed by different researchers towards the designing of LMWHs.

We synthesized a series of Ns based bio-conjugate hydrogelators NBC-1 to 4 following a previously published protocol.^25,26^ All these biocompatible NBCs showed hydrogelation property at RT and displayed highly anti-inflammatory property by reducing prostaglandin E2 (PGE2) like parental compound Ns in bacterial endotoxin lipopolysaccharide (LPS) and mouse cytokine interferon-γ (IFN-γ) induced mouse macrophage cells RAW 264.7.^25^ But, whether the molecular mechanisms behind the anti-inflammatory property of NBCs are similar to that of the parental compound Ns is still unknown. It is also unknown that the effectiveness of NBCs to anti-inflammation. We report the anti-inflammatory mechanism mediated by NBC-2, based on the inhibition of LPS/IFN-γ induced pro-inflammatory mediator NO synthesis and the reduction of the expression of inflammatory protein inducible nitric oxide synthase (iNOS) and COX-2. Reduction of iNOS expression can be correlated with the increased stability of IκBα, which in turn reduces the translocalization of transcription factor NF-κB from the cytoplasm to the nucleus. Interestingly, NBC-2 has greater binding affinity for COX-1 as compared to other NBCs. In case of LPS/IFN-γ induced inflammation, our results highlight a new insight into the underlying different molecular mechanism and correlation between the anti-inflammatory property and self-drug delivery system by potent naproxen amino alcohol bio-conjugate-2 compared with naproxen alone.

## Results

### Effect of Naproxen Bio-Conjugates on RAW 264.7 Cells

We have previously reported that the bio-conjugations of amino acid and amino-alcohol with Ns, a synthetic NSAID, were capable to produce hydrogel at a particular gelation concentration and were as biocompatible as the parent drug Ns.^25^ Table 1 shows the details of all naproxen bio-conjugates (NBC-1 to 4) including structure, molecular weight, molecular length and their gelation properties. The molecular length of NBC-1, NBC-2, NBC-3 and NBC-4, calculated by chimer software, are 29.69 Å, 21.08 Å, 20.78 Å and 20.46 Å respectively. Molecular lengths are increased by ~ 2-3 times as compared to the parental compound Ns (9.3 Å). To know the effect of these NBCs on cellular morphology, we captured images of RAW 264.7 treated with 0-3mM NBCs for 24h and showed that cells’ morphology remains unchanged in the presence of NBCs up to 0.3 mM of concentration (Supporting information, Figure S1). The cellular morphology drastically changes at 3mM concentration of NBC-1 and 2 whereas no such changes were observed with NBC-3 and 4 at same concentration. Therefore, we select 0.3mM concentration for each NBC in our subsequent experiments.

**Table 1:**
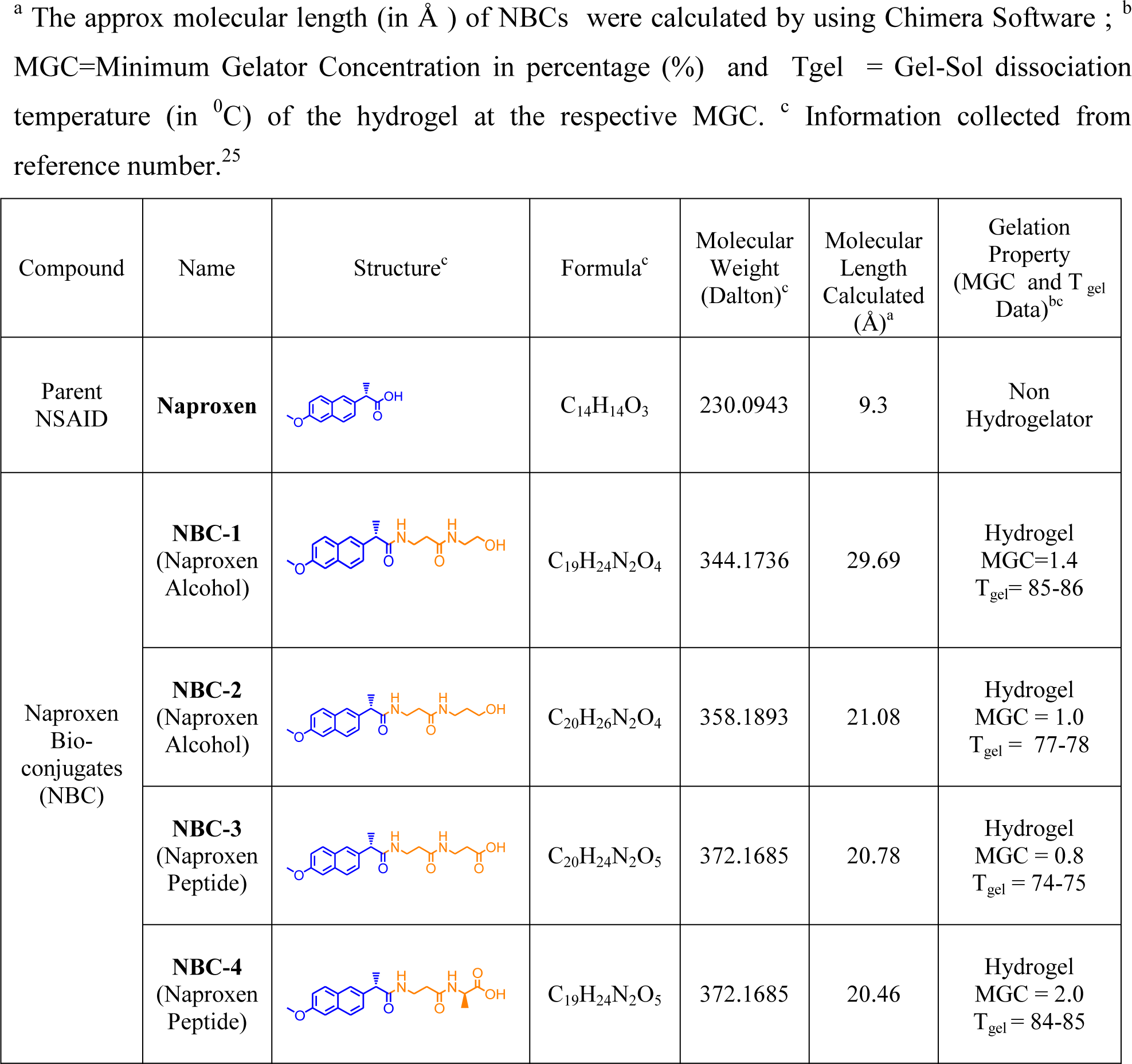
Structure and Different Parameters of Naproxen and its Bio-conjugates

<sup>a</sup> The approx molecular length (in Å) of NBCs were calculated by using Chimera Software ; <sup>b</sup> MGC=Minimum Gelator Concentration in percentage (%) and T<sub>gel</sub> = Gel-Sol dissociation temperature (in °C) of the hydrogel at the respective MGC.
| Compound | Name | Structure<sup>c</sup> | Formula<sup>c</sup> | Molecular Weight (Dalton)<sup>c</sup> | Molecular Length Calculated (Å)<sup>a</sup> | Gelation Property (MGC and T<sub>gel</sub> Data)<sup>bc</sup> |
| --- | --- | --- | --- | --- | --- | --- |
| Parent NSAID | **Naproxen** |  | C<sub>14</sub>H<sub>14</sub>O<sub>3</sub> | 230.0943 | 9.3 | Non Hydrogelator |
| Naproxen Bio-conjugates (NBC) | **NBC-1**<br>(Naproxen Alcohol) |  | C<sub>19</sub>H<sub>24</sub>N<sub>2</sub>O<sub>4</sub> | 344.1736 | 29.69 | Hydrogel<br>MGC=1.4<br>T<sub>gel</sub>= 85-86 |
|  | **NBC-2**<br>(Naproxen Alcohol) |  | C<sub>20</sub>H<sub>26</sub>N<sub>2</sub>O<sub>4</sub> | 358.1893 | 21.08 | Hydrogel<br>MGC = 1.0<br>T<sub>gel</sub> = 77-78 |
|  | **NBC-3**<br>(Naproxen Peptide) |  | C<sub>20</sub>H<sub>24</sub>N<sub>2</sub>O<sub>5</sub> | 372.1685 | 20.78 | Hydrogel<br>MGC = 0.8<br>T<sub>gel</sub> = 74-75 |
|  | **NBC-4**<br>(Naproxen Peptide) |  | C<sub>19</sub>H<sub>24</sub>N<sub>2</sub>O<sub>5</sub> | 372.1685 | 20.46 | Hydrogel<br>MGC = 2.0<br>T<sub>gel</sub> = 84-85 |

### Inhibition of LPS/IFN-γ Induced NO Synthesis and iNOS, COX-2 Expression in Presence of NBC-2

We have previously shown that LPS/IFN-γ could induce the synthesis of a potent inflammatory mediator prostaglandin E2 (PGE2), in mouse macrophage cell RAW 264.7.^25^ PGE2 is synthesized from one of the cellular membrane components, arachidonic acid (AA) by the activity of both constitutively (isoform-1) and inducibly (isoform-2) expressed cyclooxygenase enzyme (COX) (EC 1.14.99.1). We were interested to check whether LPS/IFN-γ could induce other inflammatory molecules and change the cellular morphology, a hall mark of macrophage activation.^27^ Nitric oxide (NO) is synthesized from L-arginine by both constitutive (cNOS) and inducible (iNOS) nitric oxide synthases (NOS) (EC 1.14.13.39) which are characterized as low and high NO production system respectively.^28^ During macrophage activation with LPS/IFN-γ, the iNOS gets upregulated and large amounts of NO is synthesized, considers as naturally occurring nonspecific cellular immune responses against foreign invader or tumor cells.^29^ We found that 1 μg/mL LPS and100 ng/mL IFN-γ could produce 5.75±0.32 fold increased NO synthesis compared with unstimulated cells (Supporting information, Figure S2A). Bright field images revealed that RAW 264.7 cells were more spreaded with lamellipodia like extension in presence of 1 μg/mL LPS and100 ng/mL IFN-γ (Supporting information, Figure S2B).

In order to study the anti-inflammatory properties of NBCs, LPS/IFN-γ induced inflammatory RAW264.7 cells were treated with each of NBCs and assayed for the production of NO and the expression of iNOS and COX-2. Figure 1A shows that NBCs reduce the production of NO significantly compared with Ns (p<0.0001). Note that effect of NBC-3 was relatively lesser compared with other NBCs.

**Figure 1.**
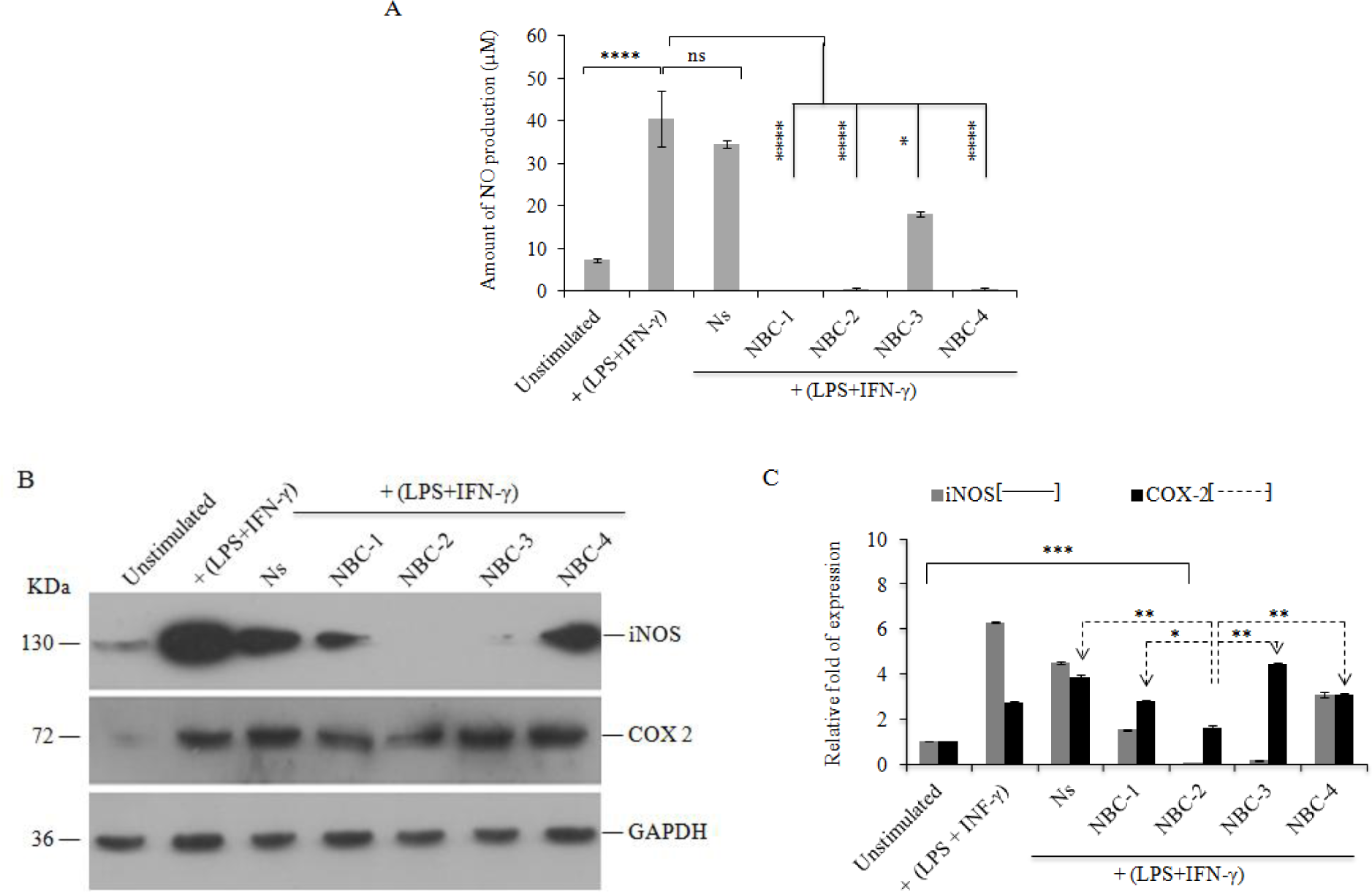
Inhibition of LPS/IFN-γ Induced NO Synthesis and iNOS, COX-2 protein expression in presence of NBC-2. RAW 246.7 cells were treated with LPS/IFN-γ and 0.3mM Ns or 0.3mM NBCs for 24h. Unstimulated and only LPS/IFN-γ treated cells were taken as –ve and +ve control respectively. (A) After incubation, conditioned cell culture media was used to measure NO production by Griess reagent. Note that, NBC-1, 2, and 4 reduce the LPS/IFN-γ induced NO synthesis than Ns and NBC-3 compared with only LPS/IFN-γ treatment. Results are means of three separate experiments (in triplicate) ± SD. (B) Immunoblot blot analysis of the expression profile of iNOS and COX-2. GAPDH was used as loading control. (C) Quantification of band intensity of iNOS and COX-2. Relative fold of expression was calculated considering the band intensity of unstimulated as “1”, indicating that NBC-2 reduces both the expression of iNOS and COX-2 compared with other NBCs. Results are means of three determinations ± SD (ns- *p*>0.05; * - *p* ≤ 0.05; ** - *p* ≤ 0.01 and *** - *p* ≤ 0.001).

In presence of LPS/IFN-γ, there are 6.31 ± 0.04 and 2.75 ± 0.02 fold increased expression of iNOS and COX-2 respectively in Raw 264.7 cells compared with unstimulated control cells, which is an indication of inducible inflammation (Figure 1B-C). Addition of Ns or any of NBCs reduces the expression of iNOS but the effect of NBC-2 and 3 was more profound in reducing iNOS expression compared with other NBCs; whereas expression of COX-2 remains unchanged in presence of Ns or any of NBCs, except NBC-2. Quantification of band intensity using ImageJ analysis revealed that Ns, NBC-1 and 4 reduce LPS/IFN-γ induced iNOS expression by 4.54 ± 0.05, 1.54 ± 0.02 and 3.12 ± 0.11 fold respectively, where as treatment with NBC-2 and 3 resulted in a 0.02 ± 0.01 (p ≤ 0.001) and 0.17 ± 0.1 (p ≤ 0.01) fold change respectively, as compared to unstimulated cells (Figure 1C). Only, NBC-2 alone reduced COX-2 expression to 1.65 ± 0.1 (p ≤ 0.001) fold as compared to unstimulated cells.

On another side, there is no such change in constitutive expression level of COX-1 after treatment with Ns and other hydrogelators (Supporting information, Figure S2C). We also investigated the expression level of the pro-inflammatory cytokine, TNF-α, in presence of Ns and NBCs in LPS/IFN-γ induced inflammatory RAW 264.7 cells. 4.22±0.22 fold increased TNF-α expression was observed in LPS/IFN-γ stimulated cells compared with unstimulated cells. Addition of NBCs reduces the expression of TNF-α more effectively than Ns 9Supporting information, Figure S3). Taken together these observation indicates that NBC-2 can reduce simultaneously the production of NO, the expression of iNOS, COX-2 and TNF-α, in LPS/IFN-γ induced inflammatory RAW 264.7 cells in a more effective manner compared with Ns and other NBCs.

### NBC-2 Alters the Stability of IκBα

Activation of toll-like receptor-4 (TLR4) by LPS/IFN-γ, followed by activation of NF-κB mediated pro-inflammatory signaling pathway is a well known phenomenon.^30,31^ On the other hand, in the LPS-stimulated RAW 264.7 cells, iNOS and COX-2 are the two pro-inflammatory gene products of the transcription factor NF-κB.^32^ Primarily, NF-κB stays in cytosol as a p50/p65 heterodimer complex with its cytosolic inhibitor protein IκBα which is again regulated by upstream protein IκB kinase (IκKα/β).^33^ One of the major mechanisms involved in the activation of NF-κB is the phosphorylation of IκBα by IκKα/β at two N-terminal serine residues due to signaling from numerous stimuli, including LPS, which ultimately leads to concomitant degradation of IκBα. It opens the nuclear localization sequence on NF-κB that allows its rapid nuclear translocation to facilitate transcription of mRNAs for pro-inflammatory proteins.^34^

In order to know whether NBCs have any effect on the stability of IκBα, we first standardize the time needed for maximum degradation of IκBα in LPS/IFN-γ induced Raw 264.7 cells and then assessed the degradation in presence of Ns or each of NBCs. Immunoblot analysis shows that LPS/IFN-γ causes degradation of IκBα within 0.5h to 1h (Supporting information, Figure S4A). Quantification revealed that 3.6 ± 0.002 (p < 0.01) and 4.9 ± 0.005 (p < 0.001) fold decrease in IκBα stability at 0.5 and 1h respectively as compared with unstimulated cells (Supporting information, Figure S4B). It also observed that the pattern of IκBα degradation and restoration is cyclically changed after each ~2-3h gap, which may due to the homeostatic model of IκBα metabolism.^35^ Simultaneously, we also measured the NO production till 24h and found that LPS/IFN-γ treated RAW 264.7 cells need 8h to reach the level of NO synthesis, observed in case of inflammation, compared with untreated cells (Supporting information, Figure S4C). This 8h time duration may be necessary to proceed the signal from the degradation of IκBα, nuclear translocation of NF-κβ, initiation of iNOS synthesis to NO production.

Surprisingly, in presence of Ns or NBCs, degradation of IκBα was inhibited and was detectable within 1h (Figure 2A). Quantification revealed that the band intensity of IκBα is 2.87 ± 0.08 (p < 0.001) fold higher in NBC-2 treated cells compared with non-treated cells. Whereas the amount of IκBα was 1.47 ± 0.08, 1.95 ± 0.09, 2.42 ± 0.1 and 2.37 ± 0.05 fold higher in Ns, NBC-1, NBC-3 and NBC-4 respectively (Figure 2B). NBC-2 increases the stability of IκBα from LPS/IFN-γ induced degradation more efficiently compared with other NBCs, which may be one of the reasons for low expression level of iNOS and COX-2 in RAW 264.7 cells (Figure 1B).

**Figure 2.**
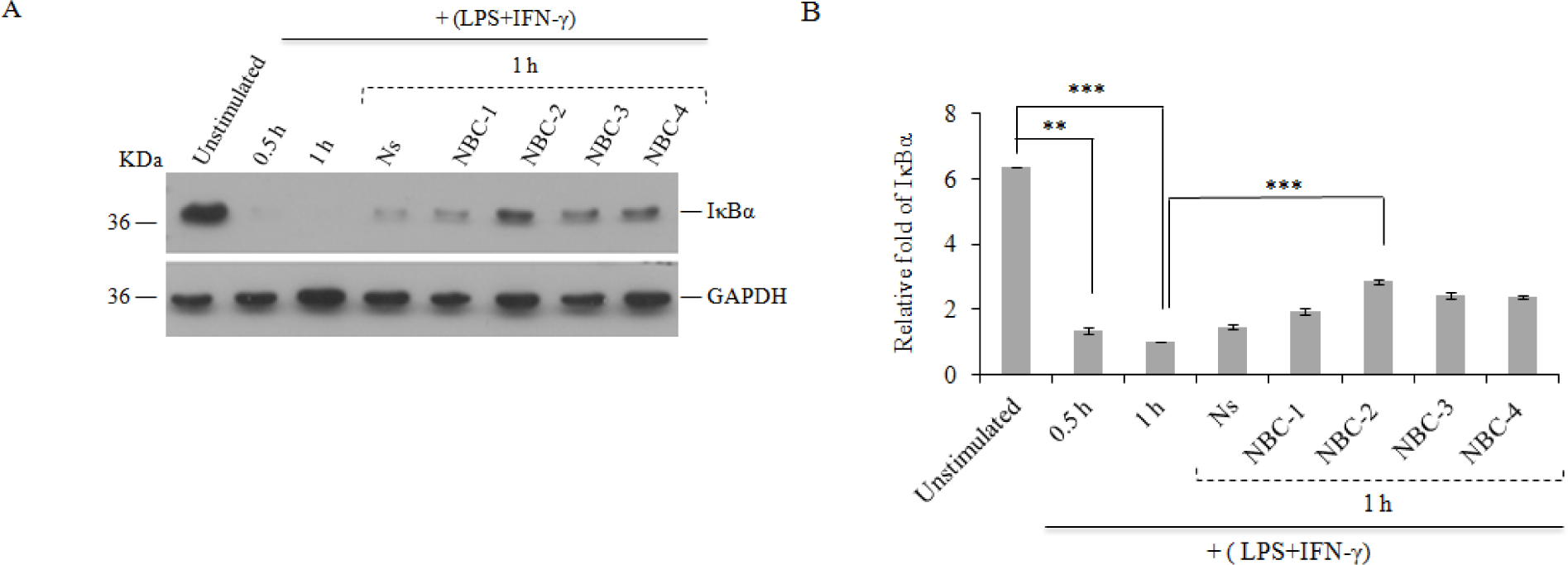
NBC-2 inhibits the LPS/IFNγ induced degradation of IκBα. RAW 246.7 cells were treated with LPS/IFN-γ and 0.3mM Ns or 0.3mM NBCs for 1h. (A) Immunoblot analysis of cell lysates with antibody against IκBα or GAPDH. GAPDH was used as loading control. Unstimulated and LPS/IFN-γ treated (0.5h and 1h) cells were taken as +ve and ‒ve control respectively. Note that IκBα was undetectable in 1h after treatment with LPS/IFN-γ; but detectable in presence of Ns or NBCs. (B) Quantification of IκBα band intensity revealed that NBC-2 has better inhibitory effect against IκBα degradation than Ns or other NBCs. Here, relative fold of IκBα expression was calculated by considering the band intensity of only LPS/IFN-γ treated RAW 264.7 cells for 1h as “1”. Results are means of three determinations ± SD (** - *p* ≤ 0.01 and *** - *p* ≤ 0.001).

### NBC-2 Inhibits LPS/IFN-γ Induced NF-κB Nuclear Translocation

A typical pro-inflammatory signaling pathway is largely based on the role of NF-κB in the expression of pro-inflammatory genes like iNOS.^36^ To evaluate the effect of NBCs on LPS/IFN-γ induced nuclear translocation of NF-κB, immunostaining of NF-κB p65 was performed with RAW 264.7 cells in the presence of each NBC (Figure 3) and was shown that inhibition of LPS/IFN-γ induced NF-κB p65 nuclear translocation by Ns and its derivatives. Image J analysis of the relative nuclear intensity of NF-κB revealed that treatment of RAW 264.7 cells with LPS/IFN-γ induces significant translocation of NF-κB p65 subunit (1.4 ± 0.6, p<0.001) to the nucleus, as compared to the unstimulated cells (0.68 ± 0.24) (Figure 3A-B). Ns (1.04 ± 0.23, p<0.05), NBC-1 (0.86 ± 0.18, p<0.01), NBC-3 (0.83 ± 0.24, p<0.01) and NBC-4 (1.06 ± 0.37, p<0.05) treated cells displayed comparatively less amount of inhibitory effect than NBC-2 (0.65 ± 0.18, p<0.001) as compared with only LPS/IFN-γ induced cells (Figure 3B); suggesting that LPS/IFN-γ induces nuclear translocation of NF-κB in RAW 264.7 cells was more effectively inhibited by NBC-2 than other NBC hydrogelators.

**Figure 3.**
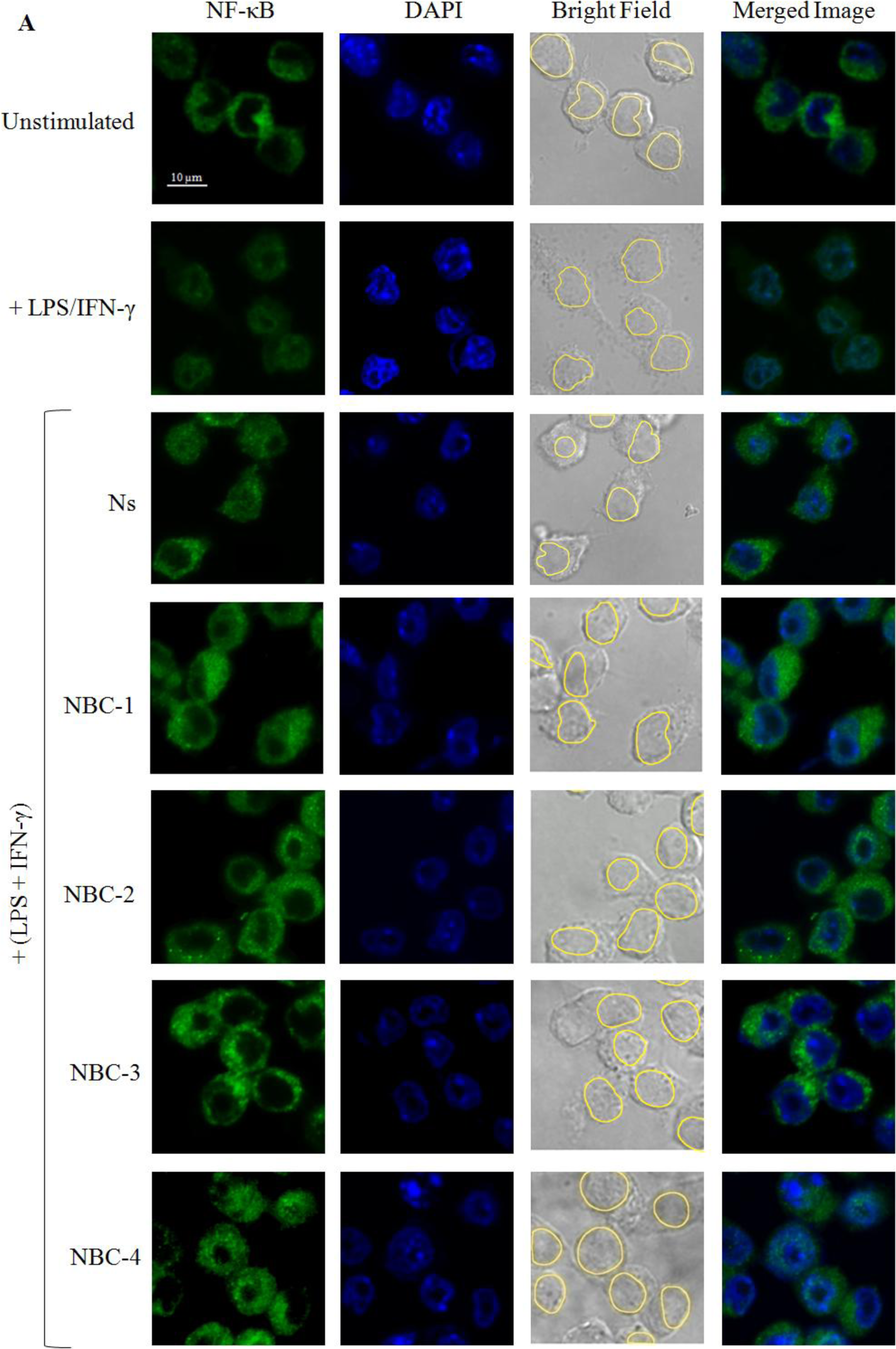
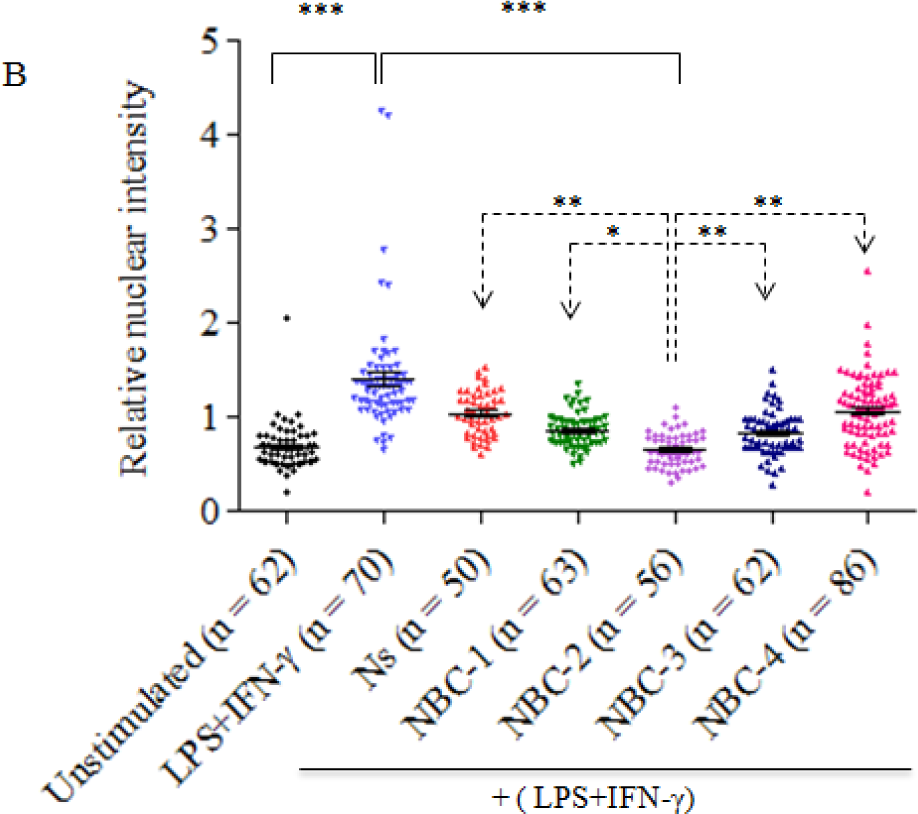
Inhibition of LPS/IFN-γ induced NF-κB nuclear translocation by NBC-2. RAW 246.7 cells were treated with LPS/IFN-γ and 0.3mM Ns or 0.3mM NBCs for 1h followed by immunostaining with NF-κBp65 antibody (green) and DAPI (blue) to visualize nucleus. (A) LPS/IFN-γ induced nuclear translocation of NF-κBp65 was determined by merged image with sea green colored. Nucleus position is marked (yellow boarder) in bright field images. Unstimulated and LPS/IFN-γ treated cells were considered as –ve and +ve control respectively. Scale bar, 10 µm. Note that reduction of NF-κB intensity in LPS/IFN-γ treated cells could be due to its translocation into the nucleus. (B) Relative nuclear intensity of NF-κB p65 was calculated by Image J analysis using a formula = [Nuclear intensity (unit/area)/whole cell intensity (unit/area)] suggesting that NBC-2 shows significant inhibitory effect compared with Ns or other derivatives [represented by dashed line (**------**)]. Result is expressed as means ± SD where “n=cell number” of different sets are mentioned into figure B. * - *P* ≤ 0.05; ** - *P* ≤ 0.01 and *** - *P* ≤ 0.001.

### Effect of Naproxen Sodium and its Derivatives on COX-1/2 Activity

Most of the NSAIDs including Ns can nonspecifically inhibit the enzyme activity of cyclooxygenase-1 (COX-1) and cyclooxygenase-2 (COX-2), followed by the synthesis inhibition of prostaglandins.^5^ To further address the question whether NBCs also have the inhibitory effect on COX-1/2 like their parent compound Ns, *in vitro* COX-1/2 enzymatic activities were determined with ovine COX-1 (oCOX-1) and human recombinant COX-2 (hCOX-2) (Figure 4). Pre-incubation of oCOX-1 and hCOX-2 with Ns, inhibited the enzyme activity in a concentration dependent manner. In presence of 300 µM Ns, the oCOX-1 and hCOX-2 activity reduced to 49.75 ± 19.06 % and 32.74 ± 1.5 % respectively (Figure 4A-B); but, there was no such effect of other NBCs except NBC-2. NBC-2 showed inhibitory effect, similar to Ns, against oCOX-1 (Figure 4A) but not against hCOX-2, suggesting that NBC-2 may have gained a selective inhibitory effect against COX-1.

**Figure 4.**
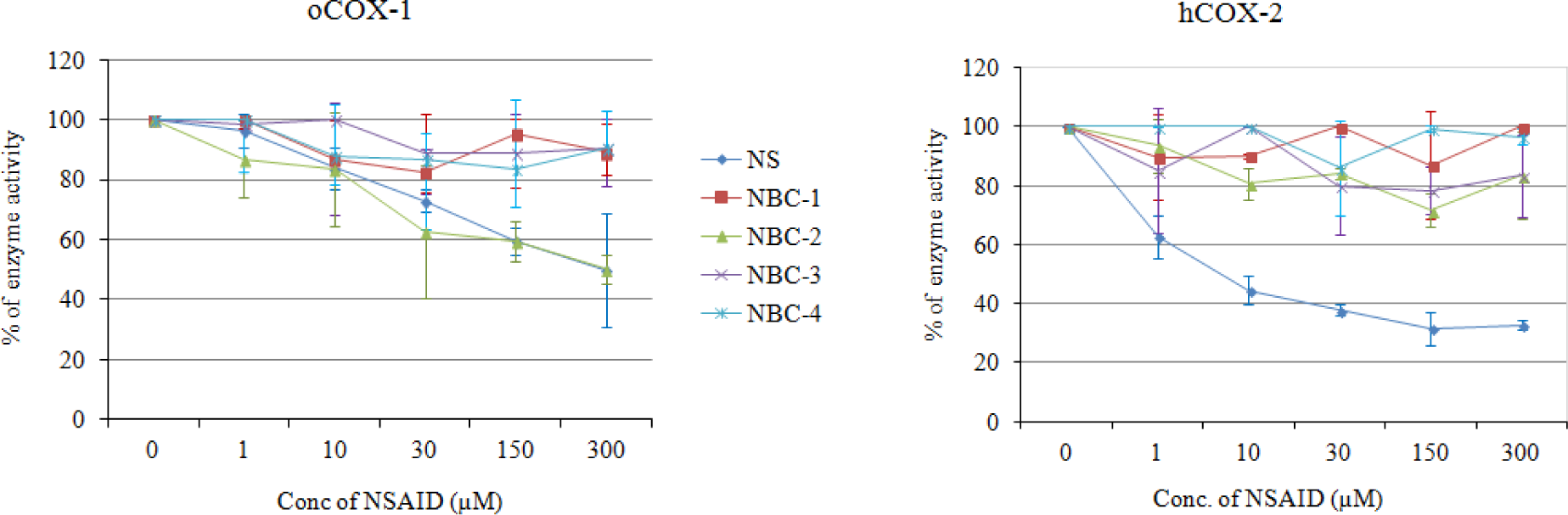
Effect of Ns and its derivatives on Cox-1/2 activity. Enzyme activity of oCOX-1 and hCOX-2 were measured after 10 min pre-incubation with different concentration (1, 10, 30, 150 and 300 µM) of Ns or NBCs at 25° C. Percent of inhibition was calculated considering the activity of enzyme in absence of compounds (0 µM) as 100%. Note that NBC-2 is more selective toward oCOX-1 rather than hCOX-2 compared with other NBCs. Results are means of three separate experiments (in triplicate) ± SD (** - *p* ≤ 0.01).

### Selectivity of NBC-2 towards COX-1

In the preceding section, we have shown in vitro that NBC-2 can preferentially inhibit ovine COX-1 (Figure 4). In order to know the molecular basis of the NBC-2 mediated inhibition of COX-1, we carried out *in silico* molecular docking analysis with Ns and NBCs by using crystal structures of COX-1 (PDB ID: 1EQG) from Ovis aris and COX-2 (PDB ID: 3NT1) from Mus musculus retrieved from the protein data bank (PDB) (Figure 5).^5,37,38^ Figure 5 shows lower COX-2 binding scores for NBC-2 derivative with respect to Naproxen. However, the difference is even more prominent in case of FlexX docking scores. Under biological conditions at pH~7, the carboxyl group of naproxen is usually deprotonated and becomes negatively charged by losing an H+.

**Figure 5.**
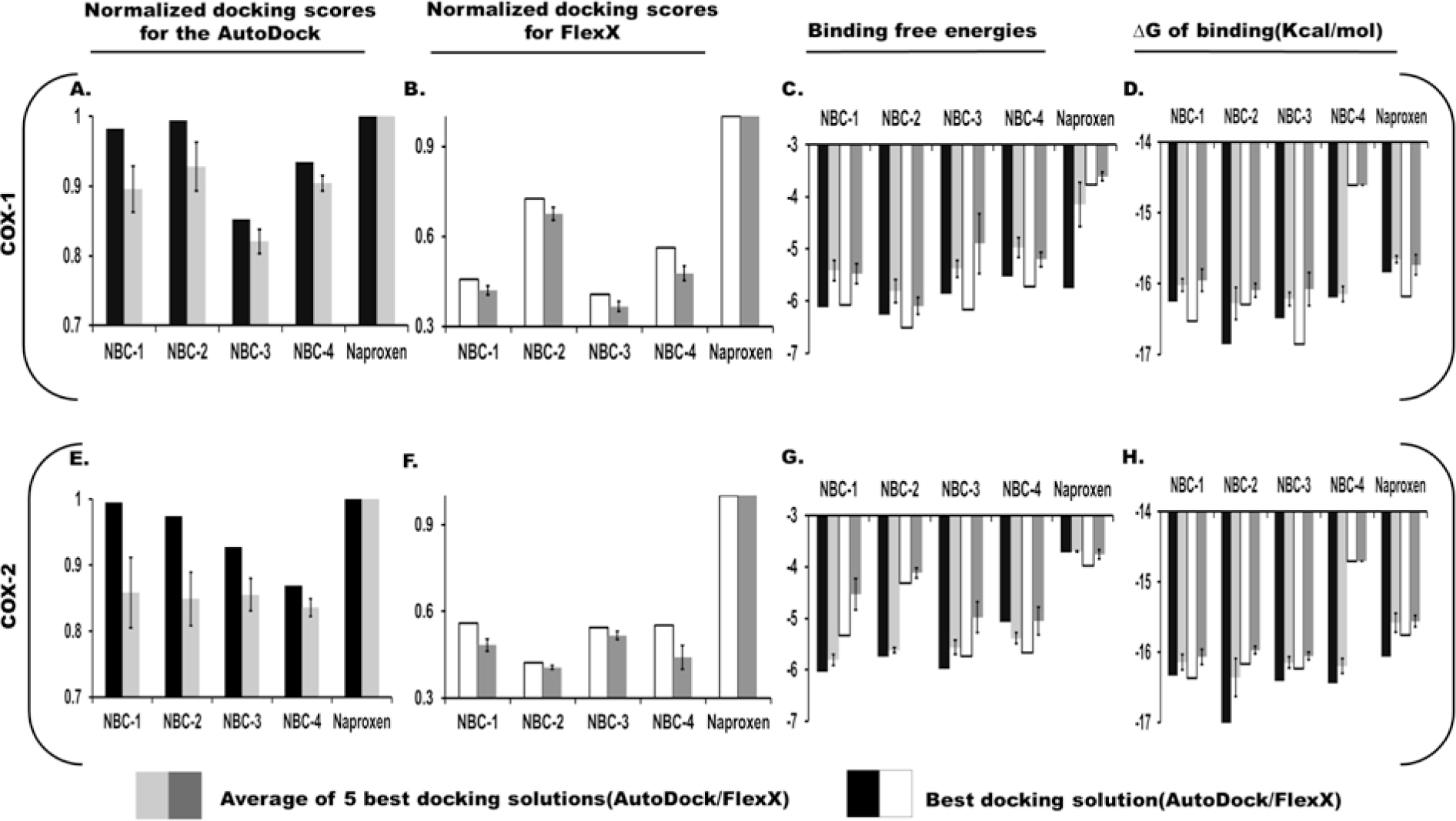
Selectivity of NBC-2 towards COX-1. Docking analysis of the Ns and its four derivatives (NBC-1, NBC-2, NBC-3, and NBC-4) with COX-1 (PDB ID: 1EQG) from *Ovis aries* and COX-2 (PDB ID: 3NT1) from *Mus musculus*. Panel A) (COX-1) and E) (COX-2) represent the normalized docking scores for the AutoDock docking solutions.^40^ Panel B) (COX-1) and F) (COX-2) represent the normalized docking scores for FlexX docking solutions.^39^ Panel C, D) (COX-1) and G, H) (COX-2) represent the respective binding free energies calculated via Cyscore^41^ and the ∆G of binding calculated via MOPAC^38^ for best and best 5 docking solutions of the COX-1 and COX-2 docking, respectively.

Carboxyl groups have an electronegative oxygen atom double bonded to a carbon atom, which increases the polarity of the bond. The electrons of the double bond migrating to another oxygen atom are stabilized by two equivalent resonance structures in which the negative charge is effectively delocalized between two more electronegative oxygen atoms of the carboxyl group of naproxen. These two electronegative atoms participate in hydrogen-bonding interactions (dipole-dipole) with the two amine group of Arg120 and the hydroxyl group of Tyr355 at the base of the active site. Moreover, the rest of naproxen inserts into the hydrophobic cleft and mostly have strong hydrophobic interactions. The p-methoxy group of naproxen is oriented toward the top of the COX-2 active site and forms van der Waals interactions with Trp387 and Tyr385 (Supporting information, Figure S5A).

In the naproxen derivative NBC-2, the oxygen atom of amide group forms H-bond interaction with Arg120 only and the terminal hydroxyl group forms H-bond interaction with Arg513 and Glu524. Although the amide group forms resonance, however it is less reactive than the carboxylate ion present in naproxen. This is due to the greater electro negativity of oxygen, the carbonyl (C=O) is a stronger dipole than the N=C dipole. There is no interaction with the hydroxyl group of Tyr355 at the base of the active site (Supporting information, Figure S5B).

The best docking orientations of the NBC-2 with COX-1 and COX-2 have been shown in figure 6A-D. Panels A and B represent the best possible docking mode of NBC-2 with COX-1 as suggested by FlexX and AutoDock, respectively.^39,40^ Both solutions suggest similar region and mode of binding. Panels C and D also show similarity in binding region and mode of NBC-2 with COX-2 between the best possible docking solutions derived from AutoDock and FlexX, respectively. Our molecular docking analysis using FlexX docking program, identify the probable interactions between NBC-2 with COX-1 and COX-2, respectively. It has been observed that NBC-2 atoms such as O26, O16, O21, H46, H52 are likely to form 7 H-bonds with COX-1 atoms, Pro86:CA, Arg120:NH2, Tyr355:OH, Glu524:OE2, respectively (Figure 6B). The complex is further stabilized by hydrophobic interactions contributed by Val349 forming Π-Sigma interaction, by Val349, Leu352, Ile523, Ala527 forming Π-Alkyl stacking and by Val116, Val349, Leu359 and Leu531 forming alkyl interactions (Supporting information, Table 1)

**Figure 6.**
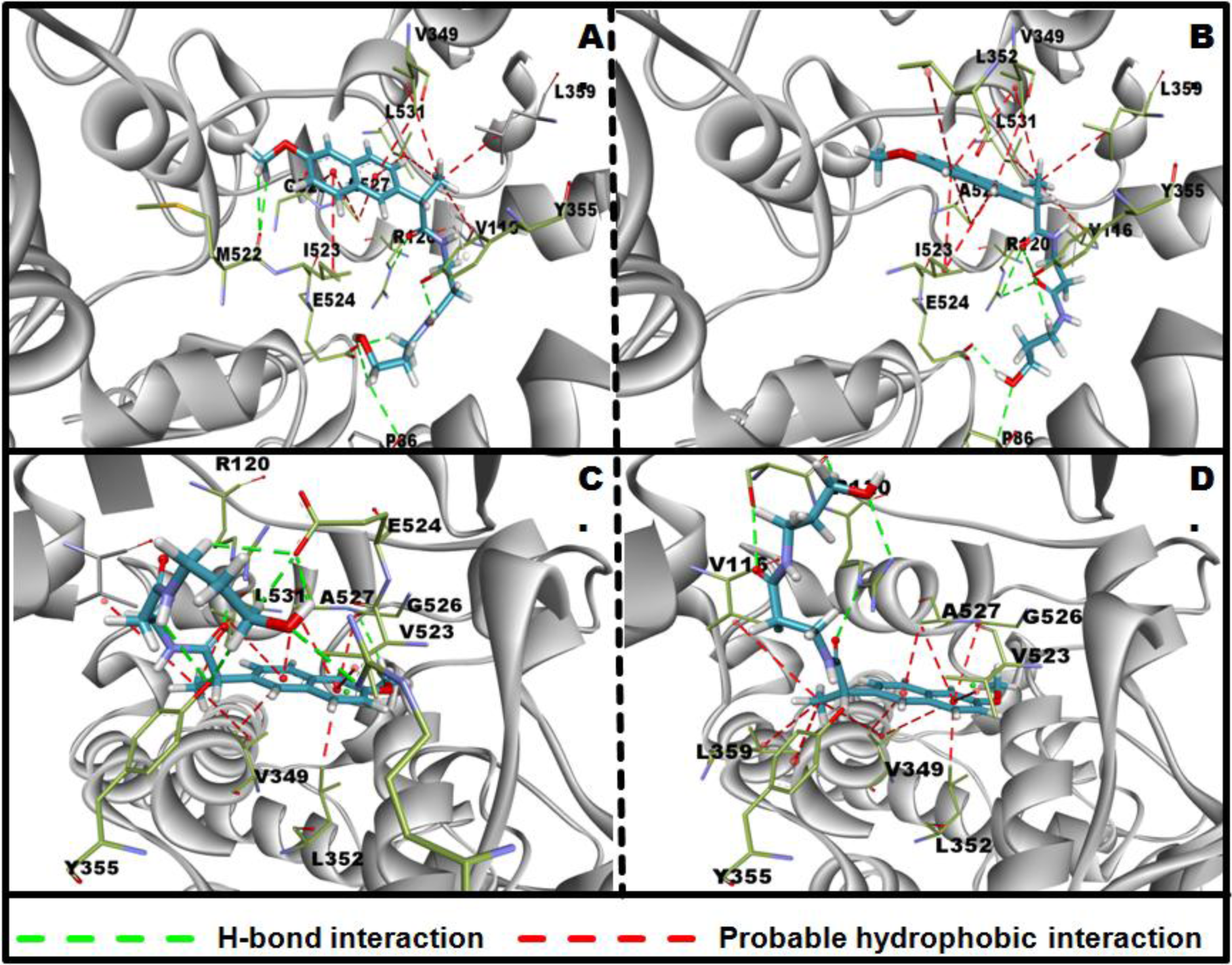
Probable docking mode of NBC-2 with COX-1 & COX-2. Docking analysis of the naproxen derivative (NBC-2) with COX-1 (PDB ID: 1EQG) from *Ovis aries* and COX-2 (PDB ID: 3NT1) from *Mus musculus*. Panel A, B) (COX-1) and panel C, D) (COX-2) represent the probable modes of binding of NBC-2 derived with COX-1 and COX-2 from AutoDock^40^ and FlexX^39^ docking solutions, respectively. Residues that are likely to be involved in hydrogen bonding and hydrophobic interaction are marked in green and red dotted lines, respectively.

On the other hand, NBC-2 atoms, O21, H47, H51, O16, O26 form 5 H-bonds with COX-2 at Ser119:OG, Ser119:O, Arg120:NE, Arg120:NH1, respectively (Fig. 6D). Probable hydrophobic interaction can also be contributed by Val349, Leu352, Ala527 forming Π-Sigma interactions, Gly526, Ala527 forming Amide-Π stacking, Tyr355, Val349, Val523, Ala527 forming Π-Alkyl interactions and Val116, Val349, Leu359 forming Alkyl interactions (Supporting information, Table 1).

Overall it has been observed that the 4 different amino acids like Pro86, Arg120, Tyr355 and Glu524 are involved in forming 7 H-bonds interactions for stabilizing the NBC-2 with COX-1 co-complex, whereas NBC-2 - COX-2 co-complex is stabilized by 5 H-bonds formed by two COX-2 protein residues, Ser119 and Arg120. Further, the interaction between top NBC-2 FlexX docked complex with COX-1 and COX-2 is compared using the Cyscore and MOPAC programs.^41,42^ The Cyscore, which is an empirical scoring function for prediction of protein-ligand binding affinity, suggests that NBC-2 has relatively higher selectivity for COX-1 over COX-2 (Figure 5C and 5G).

## Discussion

NSAIDs are among the most common pain relievers in the world and also the most controversial as well. The most effective members of this NSAID group of drugs, naproxen, aspirin and ibuprofen, are available all over the world.^43^ Naproxen has been approved by FDA (Food and Drug Administration, USA) since the mid-1970s.^5^ Due to poor water solubility of naproxen and subsequent low bioavailability,^11^ efforts have been made towards the development of naproxen based hydrogelators or naproxen bio-conjugates (NBC-1 to 4) which can form hydrogel at room temperature.^25^ Hydrogels show a sustained release of the hydrogelator (e.g. here naproxen) under physiological conditions, and this property can be used for developing a self-delivery system.

Present report provides the first evidence of an anti-inflammatory mechanism of γ-amino alcohol NBC (NBC-2) in a model cell line, mouse macrophage RAW 264.7 compared with other three NBCs. We have demonstrated that NBC-2 could be potentially used as an anti-inflammatory drug as it can reduce the expression of both LPS/IFN-γ induced pro-inflammatory proteins iNOS and COX-2 as compared with other NBCs. Though the β-amino acid NBC (NBC-3) could reduce the expression of iNOS like NBC-2 but it could not able to reduce the expression of COX-2 (Figure 1B-C). Lowering the expression of iNOS and COX-2 by NBC-2 may also lead to reduce the production of two major pro-inflammatory mediators, NO (Figure 1A) and PGE2.^25^ Since, IκBα degradation and subsequent nuclear translocation of DNA transcription factor NF-κB are the two important factors that play a critical role in the regulation of genes involved in immune response and coordinate the expression of LPS induced pro-inflammatory proteins including iNOS and COX-2,^33,35^ we have investigated the effect of NBCs on both the stability of IκBα and nuclear translocation of NF-κB. Immunoblot and immunostaining analyses revealed that NBC-2 not only significantly reduces the LPS/IFN-γ induced degradation of IκBα in cytosol but also inhibits the nuclear translocation of NF-κB (Figure 2–3) compared with other three NBCs. This effect was greater than that observed with the parental compound Ns. This observation suggests that NBC-2, but not Ns or other NBCs which are less effective than NBC-2, might perturb either phosphorylation or ubiquitination of IκBα, required for LPS/IFN-γ induced degradation of IκBα.^44,45^ This could be one of the possible mechanisms of NBC-2 mediated anti-inflammatory effect on LPS/IFN-γ treated RAW 264.7 cells. Further studies are needed to decipher the kind of modification occurred in presence of NBC-2, which inhibits the LPS/IFN-γ induced IκBα degradation in RAW 264.7 cells.

Moreover, Ns non-specifically inhibit both the COX isoforms (COX-1/2) that’s why it is quite difficult to assess isoform specific functional analysis. To gain more information on the selectivity of NBCs towards COX-1/2, in vitro enzyme assay and *in silico* molecular docking was carried out (Figure 4-5). The novelty of our study is that NBC-2 has more specificity towards COX-1, which can be useful to inhibit COX-1 selectively for analyzing isoform specific functional analysis. Addition of one -CH_2_ group creates more bending of the γ-amino alcohol group of NBC-2 in the active site of COX-1, which may provide better platform for more hydrogen bonding and hydrophobic interaction compared with β-amino alcohol group containing NBC-1 (Table 1 and Figure 6).

A single amino acid difference (V509I) in the active sites of COX-1 and 2 revealed a series of differences at the mouth of their respective active site pocket, perhaps leading to change the selectivity of specific inhibitors.^46^ Inflammation is one of the predominant features associated with most of the cancer initiation and progression.^47,48^ Recent findings have revealed that up-regulation of COX-1 expression is shown in case of human breast cancer,^49^ human prostate cancer,^50^ human Squamous cell carcinoma and adenocarcinoma,^51^ mouse lung tumor,^52^ and both human and mouse epithelial ovarian cancer.^53^ COX-1 is involved in pain processing and sensitization in spinal cord and gracile nucleus after surgery.^54^ Though, according to Grosser et al., naproxen is a more potent inhibitor of COX-2 than COX-1.^6^ A γ-amino-alcohol modification of naproxen (NBC-2) not only converts it into a potent anti-inflammatory molecule, but also increases it’s selectivity towards COX-1. The selectivity of NBC-2 for COX-1 over COX-2 may be due to loss of activity against COX-2, which may attribute to γ-amino alcohol conjugation of Ns. Although the IC_50_ of NBC-2 for COX-1 is relatively high but cells remain healthy at that IC_50_ value (Supporting information, Figure S1). Collectively, these data establish that NBC-2 can be used as a parent compound for further fine tuning to design and synthesize self-delivery anti-inflammatory drug as a hydrogel that can have lower IC_50_ value for COX-1.

## Conclusion

Taking together the results from LPS/IFN-γ induced murine macrophage RAW 264.7 model indicates that, NBC-2 shows two functions: one is to reduce the expression of COX-2 only and second to inhibit the activity of COX-1 more effectively, which is absent in other Ns derivatives. We conclude that the naproxen based hydrogelator, NBC-2 is more profound to its potential to reduce the expression of pro-inflammatory proteins (iNOS and COX-2) and pro-inflammatory mediator (NO and PGE2^25^) compared with other naproxen based modified compounds (NBC-1, 3 and 4) having hydrogelator property. The major mechanism behind NBC-2 mode of action is the increased stability of IκBα followed by reducing the nuclear translocation of DNA binding protein NF-κB (Figure 7). Additionally, NBC-2 also has selective inhibition property towards enzymatic activity of COX-1, which may provide a direction to synthesis a COX-1 specific inhibitor in future. Further *in vivo* studies are needed to evaluate its efficacy in treating inflammation.

**Figure 7.**
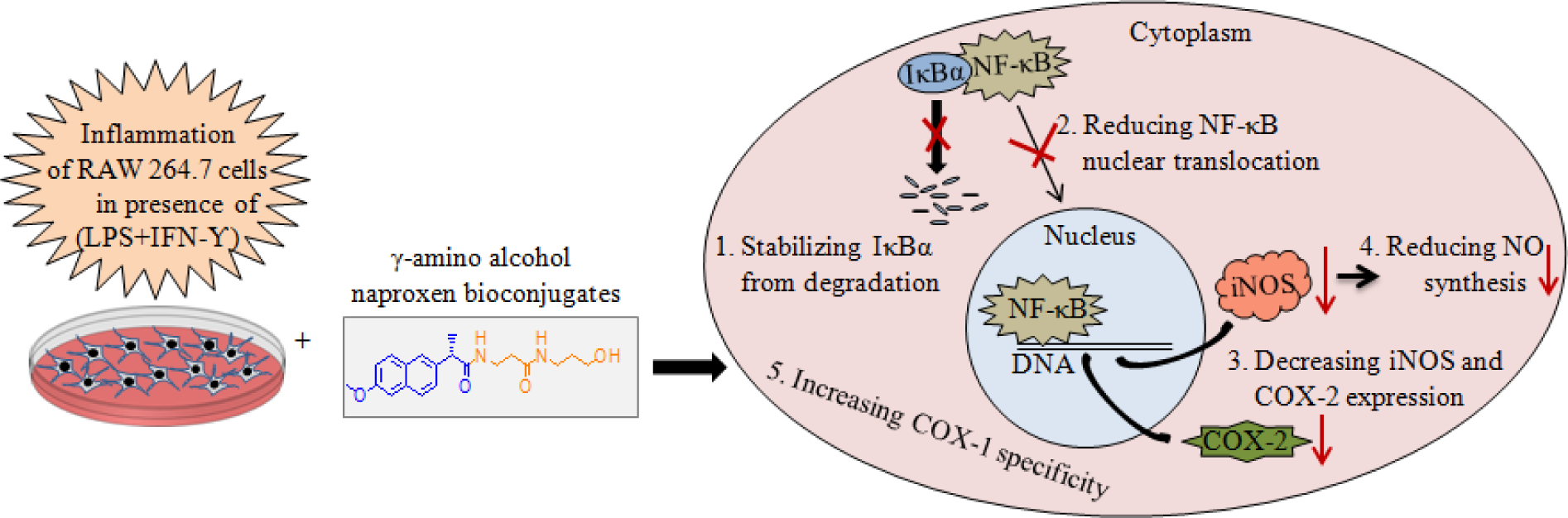
**A** model of anti-inflammatory effect of hydrogel derived from naproxen sodium in mouse macrophage RAW 264.7 cell. LPS/IFN-γ induces inflammation in murine macrophage RAW 264.7 cell. Treatment of NBC-2, which is Ns bio-conjugation with γ-amino alcohol, shows anti-inflammatory effect by stabilizing LPS/IFN-γ induce degradation of IκBα, a cytosolic inhibitory binding protein of transcription factor NF-κB; reducing nuclear translocation of DNA transcription factor NF-κB; decreasing the synthesis of pro-inflammatory factors like iNOS, COX-2, NO and PGE2^25^. Moreover, γ-amino alcohol modification of Ns also increases the selective inhibitory effect on COX-1 activity.

## Materials and methods

### Synthesis of Naproxen Bio-Conjugates

NBCs (1 to 4) were synthesized by benzotriazole coupling reaction.^25,26,55^ Firstly, Ns was conjugated with a naturally occurring β-alanine and was then further decorated with β-alanine, L-alanine and various amino-alcohols to produce the final four bio-conjugates NBC-1 to 4.

### Cell Culture for Inflammation

Mouse macrophage cell, RAW 264.7, was purchased from ATCC (Manassas, VA, USA) and maintained following their guidelines. To induce inflammation, overnight culture of 1×10^6^ RAW 264.7 cells/9.5 cm^2^, were treated with 1 μg/ml LPS from *Escherichia coli* (serotype 0111: B4, Sigma-Aldrich, St. Louis, MO, USA) and 100 ng/ml IFN-γ (ProSpec-Tany Technogene Ltd. East Brunswick, NJ, USA) for 1 or 24h. Treatment of Ns or each of NBCs was given at 0.3 mM for 1 or 24h. Cells could produce 40-80 μM NO in presence LPS/IFN-γ at 24h, which was taken as standard for the cell inflammation condition.

### Immunoblotting Analysis

Cellular protein was collected with a low-salt lysis buffer and was subjected to immunoblotting analysis with GAPDH (1:4000, Santa Cruz Biotechnology Inc., CA, USA) or iNOS (1:2000) or COX-2 (1:1000) or IkBα (1:2000), (Cell Signaling Technology, Danvers, MA, USA).^56^ Relative band intensity was quantified by using ImageJ software (National Institutes of Health, Bethesda, MD, USA) after normalizing band intensity with GAPDH.

### Nitric Oxide Assay

Nitric oxide is released from the cell into cell culture media. RAW 264.7 cells were treated with LPS/IFN-γ and Ns or NBCs for 24h. Cell culture supernatants were mixed with equal volume (1:1) of Griess reagent, incubated at RT for 10 min and the absorbance was recorded at 540 nm using a 96-well plate ELISA reader (Thermo Fisher, Rockford, IL, USA).^57^ The concentration of nitrite was calculated according to the standard curve generated from known concentrations of sodium nitrite dissolved in fresh culture media.

### Immunostaining and Immunofluorescence Microscopy

As previous report, immunostaining of RAW 264.7 cells was done by using anti-NF-κB p65 (1:500, Cell Signaling Technology) and DAPI (1:1000, Sigma) for nuclei counter staining.^58^ NF-κB intensity was quantified by ImageJ after normalizing with background. Relative nuclear intensity in dot plot was represented by following the formula=N/C; where N and C represent the nuclear and whole cell intensity (unit/area) respectively.

### In Vitro COX Enzyme Assay

Isozyme-specific inhibition study was performed by following the COX Fluorescent Inhibitor Screening Assay Kit (Cayman Chemicals, Ann Arbor, MI, USA) guidelines.^59^ Shortly, kit provided ovine COX-1 or human recombinant COX-2 were pretreated with Ns or each NBCs at 1-300 µM concentrations for 10 min in RT. The percentage of COX-1/2 activity in NBCs treated samples were calculated by considering the untreated sample as 100%.

### Molecular Docking Analysis

Crystal structures of COX-1 (PDB ID: 1EQG) co-crystallized with ibuprofen from Ovis aris and COX-2 (PDB ID: 3NT1) co-crystallized with naproxen from *Mus musculus* were retrieved from the protein data bank (PDB).^60,5,37^ Three dimensional (3D) coordinates of each naproxen derivative was generated via *in-silico* modeling using maestro from Schrödinger and the lowest energy conformer for each derivative was selected based on the local minimum conformational analysis.^61^ Molecular docking of NBCs onto the COX-1 and COX-2 structures was performed using the FlexX and AutoDock docking programs.^39,40^ Each selected docking solutions were further considered for estimation of the probable binding free energy using Cyscore and ∆G of binding using MOPAC.^41,42^ Both the tools depict the corresponding binding property and the affinity of the docked small molecule with respect to the macromolecule. Details regarding the docking protocol and scoring function are provided in the supplementary methods.

### Statistical Analysis

All immunoblots and Immunostaining data of RAW 264.7 cells are expressed as mean ± standard deviation (SD) where n=3. ELISA plate reading in case of NO estimation and COX-1/2 enzyme assay were carried out in triplicate in each time and experiment was done thrice (n=3). The two-tailed P value was calculated by unpaired *t* test where P values of <0.05 were considered statistically significant in which ns - P > 0.05; * - P ≤ 0.05; ** - P ≤ 0.01; *** - P ≤ 0.001 and ****-P≤0.0001.

## Acknowledgements

We thank Indian Association for the Cultivation of Science (IACS) and CSIR-Indian Institute of Chemical Biology (IICB) for the research work. We thank Dr. Nahid Ali (IICB) and Dr. Sib Sankar Roy (IICB) for kind gift of TNF-α antibody. We like to thank Dr. Parthasarathi Dastidar (IACS) and Dr Benu Brata Das (IACS) for helpful discussion. We also thank to Debdatta Halder, Raman Kumar Singh and Rumana Parveen for critically reading the manuscript and for other technical supports.

## Supporting Information is in separate file

## Abbreviations

NSAID: Nonsteroidal anti-inflammatory drug
Np: Naproxen
Ns: Naproxen as sodium salt form
NBC: Naproxen bio-conjugate
LPS: Lipopolysaccharide
IFN-γ: Interferon gamma
iNOS: Inducible nitric oxide synthase
COX-1 and COX-2: Cyclooxygenase enzyme isoform 1 and 2
NO: Nitric Oxide
PGE2: Prostaglandin E2
IκBα: Nuclear factor of kappa light polypeptide gene enhancer in B-cells inhibitor, alpha
NF-κB: Nuclear factor kappa-light-chain-enhancer of activated B cells
RT: Room temperature

## Footnotes

### 2. Author contributions

MRD designed and conducted all studies with the help of other co-authors. MRD and SSJ conceived the study and analyzed all data except *in silico* molecular docking analysis which was carried out by AB and SC. Compounds were synthesized by JM. MRD, AB, SC, and SSJ wrote the manuscript. All the authors have read and approved the final manuscript.

