## supplemental figure for "Amino-alcohol Bio-conjugate of Naproxen Exhibits Anti-inflammatory Activity through NF-κB Signaling Pathway"

### Author Information

[§]Department of Biological Chemistry; Indian Association for the Cultivation of Science; 2A & 2B, Raja Subodh Chandra Mallick Road; Jadavpur, Kolkata, West Bengal 700032, India.

[†]Present address: Department of Biology of Cancer & Other Diseases; National Centre for Cell Science; NCCS Complex; University of Pune Campus; University Road, Ganeshkhind, Pune, Maharashtra 411007, India

[¶]Structural Biology and Bio-informatics Division; CSIR-Indian Institute of Chemical Biology; 4, Raja Subodh Chandra Mallick Road; Poddar Nagar; Jadavpur, Kolkata, West Bengal 700032, India.

[††]Present address: Department of Bioinformatics, Tata Translational Cancer Research Centre, Tata Medical Center, Kolkata-700156, India.

[§§] Director's Research Unit, Indian Association for the Cultivation of Science; 2A & 2B, Raja Subodh Chandra Mallick Road; Jadavpur, Kolkata, West Bengal 700032, India.

[‖] Department of Organic Chemistry, Indian Association for the Cultivation of Science; 2A & 2B, Raja Subodh Chandra Mallick Road; Jadavpur, Kolkata, West Bengal 700032, India.

[#]Present address: Department of Chemistry, Purdue University Centre for Drug Discovery, Purdue University, West Lafayette, IN 47906, USA.

### Supplementary Materials and methods

#### Effect of Naproxen Bio-conjugates on Morphology of RAW 264.7 Cells

Effect of naproxen bio-conjugates (NBC-1 to 4) on the phenotype of RAW 264.7 cells was screened by bright field microscopy. $1\times10^6$ RAW 264.7 cells/9.5 $cm^2$ were seeded and grown overnight in complete DMEM (supplemented with 10% FBS, 100 units/ml Penicillin, 100 µg/ml Streptomycin and 0.292 mg/ ml L-Glutamine) at 37 °C in a humidified incubator with an atmosphere of 5% CO2. Cells were then treated with different concentrations (0.00003, 0.0003, 0.03, 0.3 and 3 mM) of NBCs and incubated for 24h. Images of each well were captured by using an Olympus IX-51 microscope at 10X magnification. Scale bar, 50 µm.

#### Nitric Oxide Assay

Induction of Inflammation in RAW 264.7 cells was detected by measuring the nitric oxide (NO) production (see the "Materials and methods" in main paper) in conditioned media at 24h after treatment with different concentration of LPS/IFN-γ (1 µg/ml LPS, 1 µg/ml LPS + 30 ng/ml IFN-γ, and 1 µg/ml LPS + 100 ng/ml IFN-γ). Cells treated with media only were taken as control. Fold of NO production was calculated by considering amount of NO produced by control cells as "1". In case of time dependent NO measurement; cells were treated with LPS/IFN-γ for different time points, like 0.5, 1, 2, 4, 6, 8, 10 and 24h.

#### Inflammation in LPS/IFN-γ induced RAW 264.7 Cells

RAW 264.7 cells treated with different concentration of LPS/IFN-γ (1 µg/ml LPS, 1 µg/ml LPS + 30 ng/ml IFN-γ, and 1 µg/ml LPS + 100 ng/ml IFN-γ) were incubated for 24 h. Images RAW 264.7 cells morphology in each condition were captured by using an Olympus IX-51 microscope at 40X magnification. Scale bar, 50 µm.

### Electrophoresis and Immunoblotting

RAW 264.7 cells were treated with LPS/IFN-γ and Ns or NBCs for 24h and then were subjected to SDS-PAGE followed by immunoblot analysis (see the "Materials and methods" in main paper) by using primary antibodies to GAPDH (1:4000) or COX-1 (1:1000, Cell Signaling Technology, Danvers, MA, USA) or TNF-α (1:1000, Santa Cruz Biotechnology Inc., CA, USA). In case of IκBα, treatment of cells with only LPS/IFN-γ was carried out for different time points, like 0.5, 1, 2, 4, 6, 8 and 10h. Unstimulated cells at 10h, was considered as control.

### Molecular docking protocol

Crystal structures of COX-1 (PDB ID: 1EQG) co-crystallized with ibuprofen from Ovis aris and COX-2 (PDB ID: 3NT1) co-crystallized with naproxen from Mus musculus were retrieved from the protein data bank (PDB).[1-3] Structural comparison analysis revealed that the residues contributing to the binding site of NSAID remains structurally and sequentially conserved in both the cyclooxygenase structures with overall sequence identity of 100% and structural root mean square deviation (RMSD) of 0.74Å. Three dimensional (3D) coordinates of each naproxen derivative was generated via in-silico modeling using maestro from Schrödinger34 and the lowest energy conformer for each derivative was selected based on the local minimum conformational analysis. Molecular docking of NBCs onto the COX-1 and COX-2 structures was performed using the FlexX and AutoDock docking programs.[4-5] FlexX provides a flexible docking protocol that uses incremental construction algorithm to place the ligand into the active site of the macromolecule whereas AutoDock provides the rigid-flexible docking solutions, where it retains the rigidity for the macromolecule but provides flexibility to the ligand. Default parameters were used for FlexX. For AutoDock, protein and ligand structures were prepared by adding hydrogens and assigning gastegier charges. AutoTors utility was used to specify the tensional degrees of freedom for the ligand. Autogrid was used for grid map settings. The grids were prepared with 80x80x80Å and 86x86x86Å cube for respective COX-1 and COX-2 structures and the spacing was set at 0.375Å for both. Lamarckian genetic algorithm was used in docking simulation, with an initial population of 150 randomly placed individuals, maximum number of 2, 50,000 energy evaluations, 150000 generations, mutation rate of 0.02, a cross over rate of 0.8 and an elitism value of 1 were used. 50 docking runs were performed. Pseudo-Solis and Wets algorithm were used for local search method. Finally, the resulting docked conformations were clustered together on the basis of root mean square deviation (RMSD) tolerance of 1.5 Å and represented by most favorable free energy of binding.

To standardize the docking protocol and to identify the probable docking orientation of the naproxen derivatives with respect to the COX-1 and COX-2 crystal structures, we initially docked ibuprofen and naproxen onto the COX-1 and COX-2 structure using FlexX and AutoDock. All the top docked solutions showed high similarities in docking orientation and binding energy calculations with respect to the crystal structures of respective COX-1 and COX-2 (data not shown). Next, the naproxen and its derivatives, NBC-1, NBC-2, NBC-3 and NBC-4 were docked with corresponding COX-1 and COX-2 crystal structures. In each docking study, the best docking solution and best 5 docking solutions were compared with the top naproxen docking score with respect to the COX-1 and COX-2 and the normalized docking scores were calculated using the following formula:

$$DS_{norm} = 1 - \left(\frac{BDS_{npx} - BDS_{npxDv}}{BDS_{npx}}\right)$$

Where, DSnorm is the normalized docking score for a particular naproxen derivative with respect to a particular cyclooxygenase structure. BDSnpx and BDSnpxDV are the scores of the best docked solutions of naproxen and its derivative. Similarly, the average normalized docking score of the top 5 docking solutions were calculated by simply taking the average of DSnorm.

### Supplementary Figure Legends

**Figure S1.** Effect of NBCs on morphology of RAW 264.7 cells. RAW 264.7 cells were treated with different concentrations (0.00003, 0.0003, 0.03, 0.3 and 3 mM) of NBCs for 24 h. Bright field images of each well were captured by using 10X magnification. Scale bar, 50 µm. Note that, in case of all NBCs, there is no such change in cellular morphology up to 0.3 mM concentration. But, at 3 mM concentration, cells treated with NBC-1 and 2 show a change in morphology.

**Figure S2.** Inflammation in RAW 264.7 cells by LPS/IFN-γ. RAW 264.7 cells were treated with different concentration of LPS/IFN-γ (1 µg/ml LPS, 1 µg/ml LPS + 30 ng/ml IFN-γ and 1 µg/ml LPS + 100 ng/ml IFN-γ) for 24 h. A) NO production was measured from conditioned media by Griess reagent. Unstimulated cells were taken as control. Fold was calculated by considering amount of NO produced by control cells as “1”. Note that, 1 µg/ml LPS and 100 ng/ml IFN-γ induced NO synthesis is 5.75 ± 0.32 fold higher than compared with control. Results are means of three determinations (in triplicate) ± SD (*** - $p \leq 0.001$). B) Bright field microscopic images under same treatment condition of RAW 264.7 cells. Images of each well were captured by using 40X magnification. Scale bar, 50 μm. Note that, there is an increased number of filopodia, lamellipodia (yellow arrow) and more stretch cellular form in presence of 1 μg/ml LPS and100 ng/ml IFN-γ compared with control. C) After 24 h incubation of Ns or NBCs along with LPS/IFN-γ, expression profile of COX-1 was determined by immunoblot analysis with antibodies against COX-1 or GAPDH. GAPDH was used as loading control.

**Figure S3.** Expression of TNF-α in the presence of NBC-2. Inflammatory RAW 246.7 Cells were treated 0.3mM Ns or 0.3mM NBCs. At 24h, expression of TNF-α was determined by immunoblot analysis A) with antibody against TNF-α. GAPDH was used as loading control. B) Quantification of band intensity of TNF-α. Relative fold of expression was calculated considering the band intensity of unstimulated as "1". Note that NBC-2 (*lane 5*) reduces the expression level of TNF-α more profoundly than other NBCs. Results are means of three determinations ± SD (** - $p \leq 0.01$ and *** - $p \leq 0.001$).

**Figure S4.** Time dependent IκBα expression. RAW 264.7 cells were treated with LPS/IFN-γ for different time points (0.5, 1, 2, 4, 6, 8 and 10 h). A) Immunoblot analysis was done by using anti-IκBα and anti-GAPDH antibody. GAPDH was used as loading control. Unstimulated cells were taken as +ve control. B) Quantification of relative band intensity revealed that expression of IκBα in 0.5 h and 1 h LPS/IFN-γ treated RAW 264.7 cells are less compared with control one. Fold was calculated by considering normalized band intensity of control cells as "1". Results are means of three determinations (in triplicate) ± SD (** - $p \leq 0.01$ and *** - $p \leq 0.001$). C) At the same time point, measuring of NO with conditioned media indicates that NO production was increasing gradually from 8h (black arrow).

**Figure S5.** Interaction analysis between naproxen and its derivative NBC-2 with COX-2 (PDB ID: 3NT1). Panel A and panel B represents the probable mode of binding of naproxen and its derivative (molecule 2) with COX-2 (PDB ID: 3NT1). Image was created using LIGPLOT.[1]

**Supporting Table-1**: List of probable interaction types between the NBC- 2 with COX-1 and COX-2 respectively.

| Ligand | Protein & Docking program | Protein residue | Ligand residue | Bond type | Interaction type | Distance (Å) |
|---|---|---|---|---|---|---|
| NBC-2 | COX-1 (AutoDock) | PRO86:O | H22 | Hydrogen Bond | Conventional Hydrogen Bond | 2.81 |
| | | ARG120:NE | O16 | Hydrogen Bond | Conventional Hydrogen Bond | 2.74 |
| | | TYR355:OH | H45 | Hydrogen Bond | Conventional Hydrogen Bond | 2.02 |
| | | GLU524:OE2 | H52 | Hydrogen Bond | Conventional Hydrogen Bond | 1.88 |
| | | GLU524:OE2 | H22 | Hydrogen Bond | Carbon Hydrogen Bond | 2.67 |
| | | GLU524:OE2 | H19 | Hydrogen Bond | Carbon Hydrogen Bond | 2.13 |
| | | MET522:O | H25 | Hydrogen Bond | Carbon Hydrogen Bond | 2.91 |
| | | MET522:O | H26 | Hydrogen Bond | Carbon Hydrogen Bond | 2.64 |
| | | VAL349:CG1 | LIGAND | Hydrophobic | Pi-Sigma | 3.87 |
| | | ALA527:CB | LIGAND | Hydrophobic | Pi-Sigma | 3.40 |
| | | GLY526:C,O | LIGAND | Hydrophobic | Amide-Pi Stacked | 4.16 |
| | | ALA527:N | LIGAND | Hydrophobic | Amide-Pi Stacked | 4.16 |
| | | LEU531 | LIGAND | Hydrophobic | Pi-Alkyl | 5.24 |
| | | ILE523 | LIGAND | Hydrophobic | Pi-Alkyl | 4.80 |
| | | VAL116 | LIGAND | Hydrophobic | Alkyl | 4.78 |
| | | VAL349 | LIGAND | Hydrophobic | Alkyl | 4.90 |
| | | LEU359 | LIGAND | Hydrophobic | Alkyl | 4.11 |
| | | LEU531 | LIGAND | Hydrophobic | Alkyl | 5.03 |
| | | ALA527 | LIGAND | Hydrophobic | Pi-Alkyl | 4.19 |
| NBC-2 | COX-1 | PRO86:CA | O26 | Hydrogen Bond | Carbon Hydrogen Bond | 2.71 |

| | | | | | | |
|---|---|---|---|---|---|---|
| | (FlexX) | ARG120:NH2 | O16 | Hydrogen Bond | Conventional Hydrogen Bond | 2.96 |
| | | ARG120:NH2 | O21 | Hydrogen Bond | Conventional Hydrogen Bond | 2.65 |
| | | TYR355:OH | O16 | Hydrogen Bond | Conventional Hydrogen Bond | 3.11 |
| | | TYR355:OH | O21 | Hydrogen Bond | Conventional Hydrogen Bond | 2.41 |
| | | TYR355:OH | H46 | Hydrogen Bond | Carbon Hydrogen Bond | 2.19 |
| | | GLU524:OE2 | H52 | Hydrogen Bond | Conventional Hydrogen Bond | 2.12 |
| | | VAL349:CG1 | LIGAND | Hydrophobic | Pi-Sigma | 3.88 |
| | | VAL116 | LIGAND | Hydrophobic | Alkyl | 4.83 |
| | | VAL349 | LIGAND | Hydrophobic | Alkyl | 4.91 |
| | | VAL349 | LIGAND | Hydrophobic | Pi-Alkyl | 5.28 |
| | | LEU359 | LIGAND | Hydrophobic | Alkyl | 4.86 |
| | | LEU531 | LIGAND | Hydrophobic | Alkyl | 4.35 |
| | | LEU352 | LIGAND | Hydrophobic | Pi-Alkyl | 5.06 |
| | | ILE523 | LIGAND | Hydrophobic | Pi-Alkyl | 4.56 |
| | | ILE523 | LIGAND | Hydrophobic | Pi-Alkyl | 4.51 |
| | | ALA527 | LIGAND | Hydrophobic | Pi-Alkyl | 4.96 |
| | | ALA527 | LIGAND | Hydrophobic | Pi-Alkyl | 4.76 |
| NBC-2 | COX-2 (AutoDock) | ARG120:NE | O16 | Hydrogen Bond | Conventional Hydrogen Bond | 2.80 |
| | | ARG513:NH1 | O26 | Hydrogen Bond | Conventional Hydrogen Bond | 2.98 |
| | | TYR355:OH | H45 | Hydrogen Bond | Conventional Hydrogen Bond | 2.88 |
| | | TYR355:OH | H22 | Hydrogen Bond | Carbon Hydrogen Bond | 2.41 |
| | | GLU524:OE2 | H52 | Hydrogen Bond | Conventional Hydrogen Bond | 2.16 |
| | | GLU524:OE2 | H19 | Hydrogen Bond | Conventional Hydrogen Bond | 2.87 |
| | | GLU524:OE2 | H23 | Hydrogen Bond | Conventional Hydrogen Bond | 2.17 |
| | | GLY526:O | H25 | Hydrogen Bond | Conventional Hydrogen Bond | 2.58 |

| | | | | | | |
|---|---|---|---|---|---|---|
| | | VAL349:CG1 | LIGAND | Hydrophobic | Pi-Sigma | 3.97 |
| | | LEU352:CD1 | LIGAND | Hydrophobic | Pi-Sigma | 3.65 |
| | | ALA527:CB | LIGAND | Hydrophobic | Pi-Sigma | 3.38 |
| | | ALA527 | LIGAND | Hydrophobic | Pi-Alkyl | 4.24 |
| | | ALA527:N | LIGAND | Hydrophobic | Amide-Pi Stacked | 4.03 |
| | | VAL116 | LIGAND | Hydrophobic | Alkyl | 5.34 |
| | | VAL349 | LIGAND | Hydrophobic | Alkyl | 4.47 |
| | | LEU531 | LIGAND | Hydrophobic | Alkyl | 4.93 |
| | | LEU531 | LIGAND | Hydrophobic | Pi-Alkyl | 5.34 |
| | | VAL523 | LIGAND | Hydrophobic | Pi-Alkyl | 5.19 |
| NBC-2 | COX-2 (FlexX) | SER119:OG | O21 | Hydrogen Bond | Conventional Hydrogen Bond | 2.90 |
| | | SER119:OG | H47 | Hydrogen Bond | Carbon Hydrogen Bond | 2.91 |
| | | SER119:O | H51 | Hydrogen Bond | Carbon Hydrogen Bond | 2.23 |
| | | ARG120:NE | O16 | Hydrogen Bond | Conventional Hydrogen Bond | 2.85 |
| | | ARG120:NH1 | O26 | Hydrogen Bond | Conventional Hydrogen Bond | 2.72 |
| | | VAL349:CG1 | LIGAND | Hydrophobic | Pi-Sigma | 3.82 |
| | | LEU352:CD1 | LIGAND | Hydrophobic | Pi-Sigma | 3.32 |
| | | ALA527:CB | LIGAND | Hydrophobic | Pi-Sigma | 3.61 |
| | | GLY526:C,O | LIGAND | Hydrophobic | Amide-Pi Stacked | 4.43 |
| | | ALA527:N | LIGAND | Hydrophobic | Amide-Pi Stacked | 4.43 |
| | | VAL116 | LIGAND | Hydrophobic | Alkyl | 5.06 |
| | | VAL349 | LIGAND | Hydrophobic | Alkyl | 5.38 |
| | | LEU359 | LIGAND | Hydrophobic | Alkyl | 4.28 |
| | | TYR355 | LIGAND | Hydrophobic | Pi-Alkyl | 4.46 |
| | | VAL349 | LIGAND | Hydrophobic | Pi-Alkyl | 5.42 |

| | | | | | | |
|---|---|---|---|---|---|---|
| | | VAL523 | LIGAND | Hydrophobic | Pi-Alkyl | 5.16 |
| | | ALA527 | LIGAND | Hydrophobic | Pi-Alkyl | 4.29 |

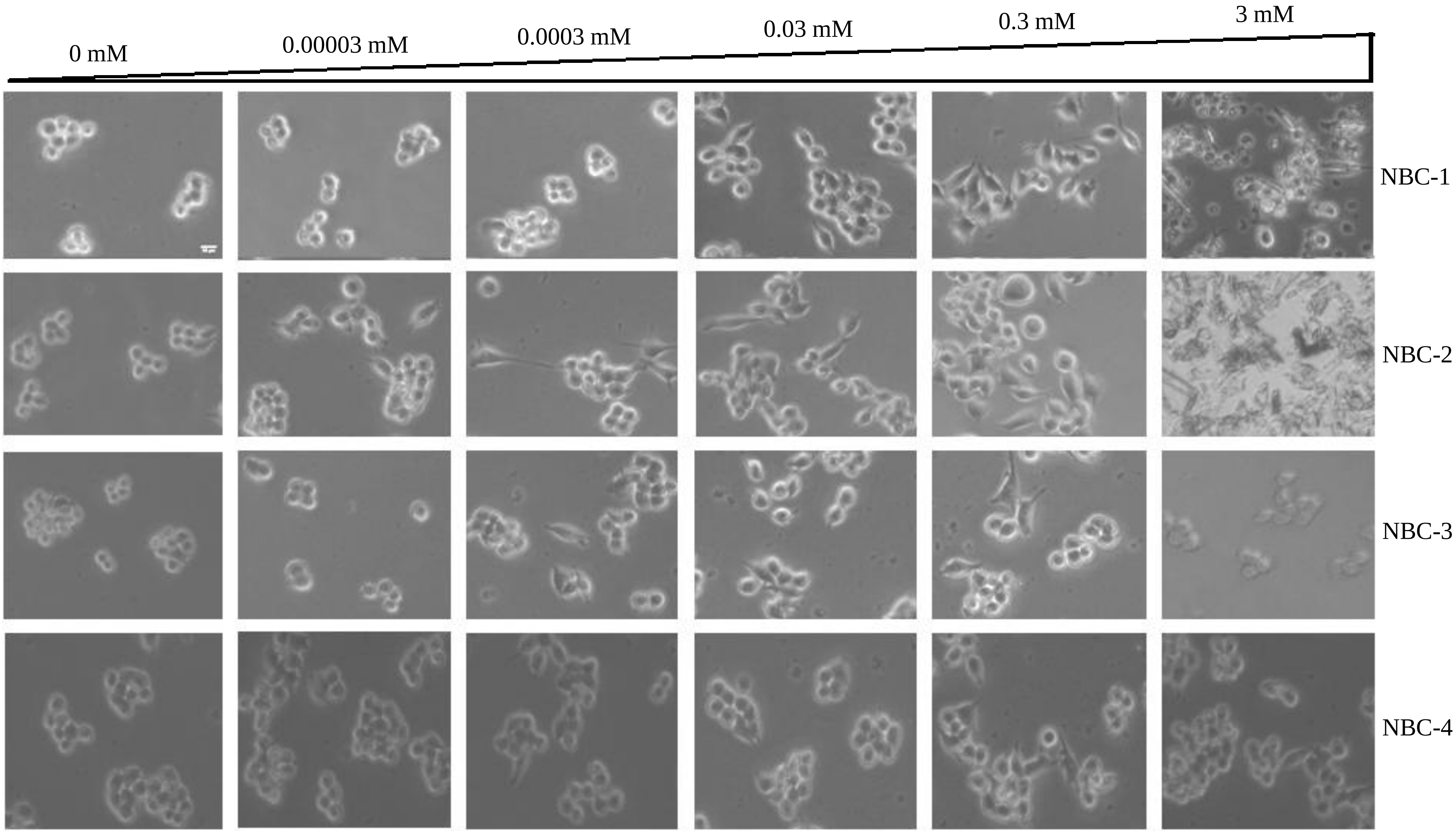

Figure S1

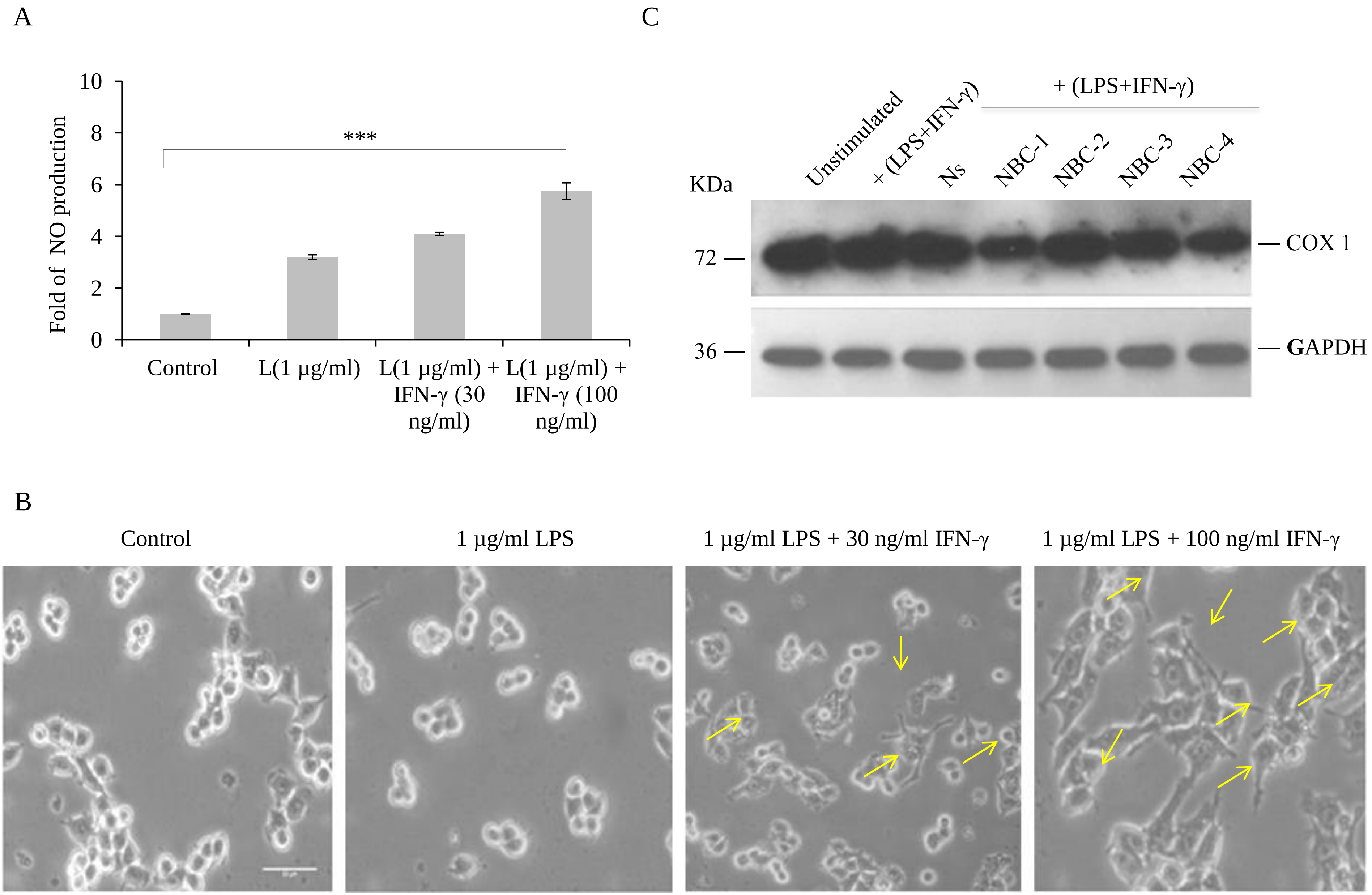
A
Fold of NO production
10
8
6
4
2
0
***
Control
L(1 µg/ml)
L(1 µg/ml) + IFN-γ (30 ng/ml)
L(1 µg/ml) + IFN-γ (100 ng/ml)
C
Unstimulated
+ (LPS+IFN-γ)
Ns
+ (LPS+IFN-γ)
NBC-1
NBC-2
NBC-3
NBC-4
KDa
72
36
COX 1
GAPDH
B
Control
1 µg/ml LPS
1 µg/ml LPS + 30 ng/ml IFN-γ
1 µg/ml LPS + 100 ng/ml IFN-γ

Figure S2

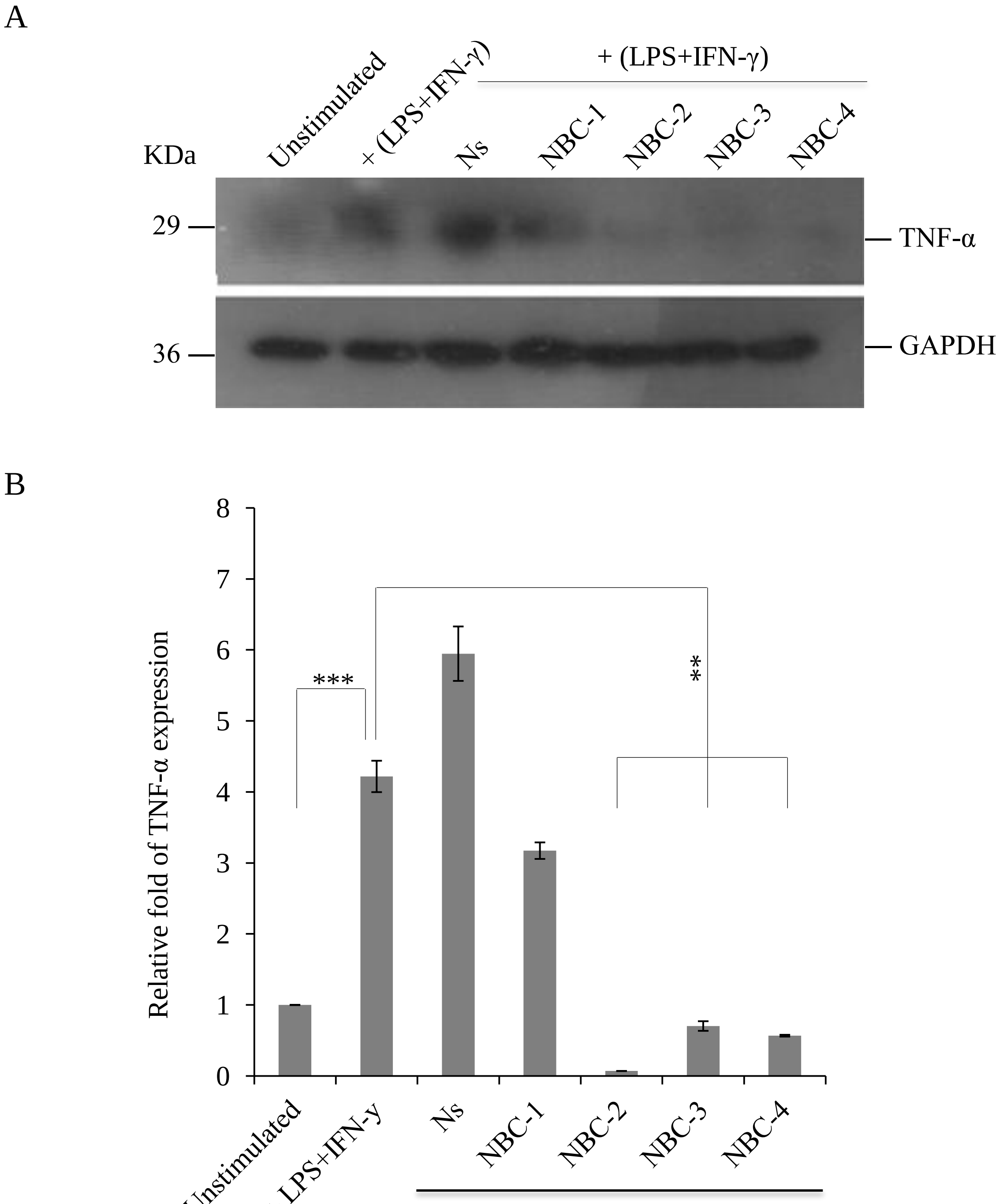

S13

Figure S3

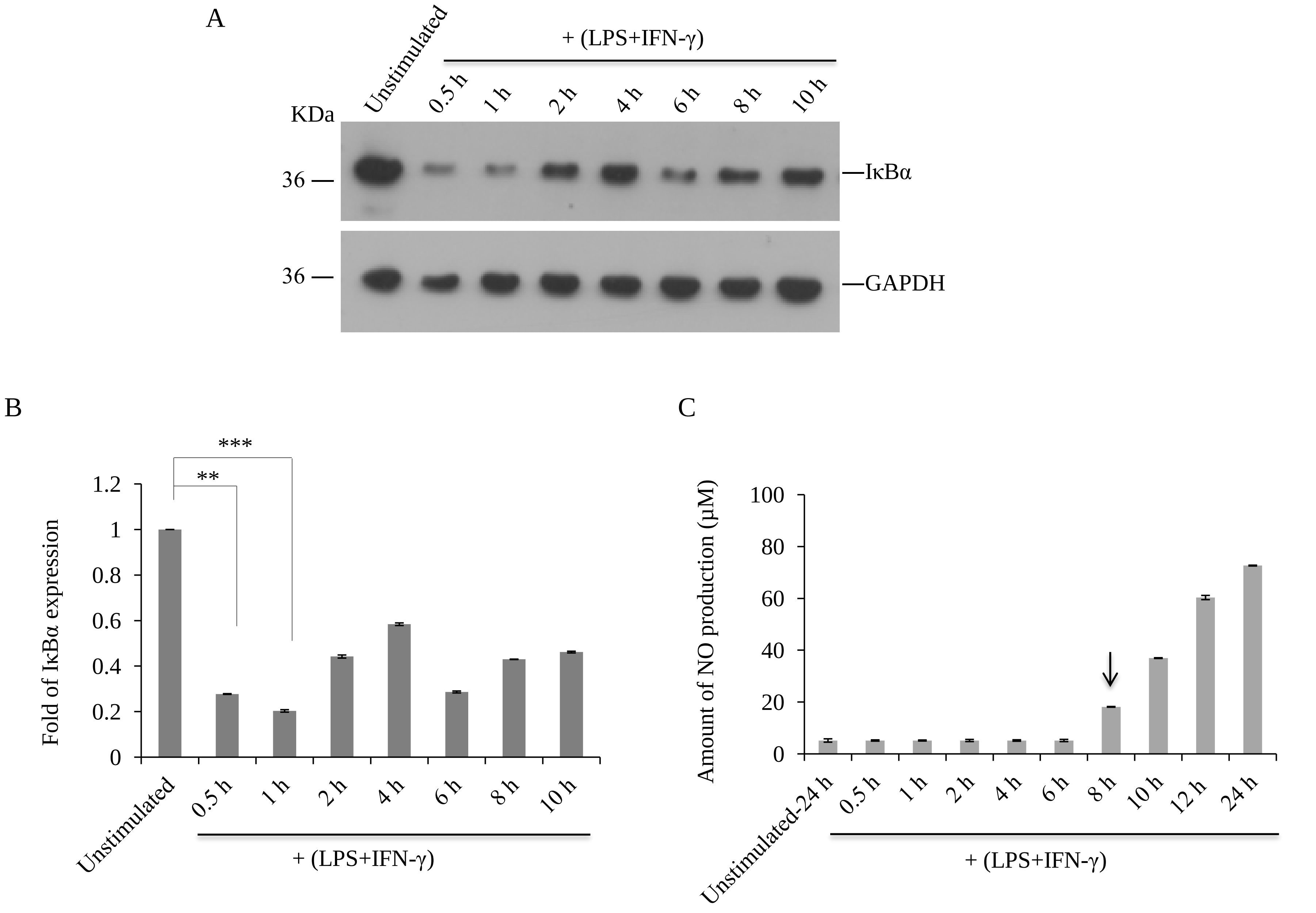
A
Unstimulated
+ (LPS+IFN-γ)
0.5 h
1 h
2 h
4 h
6 h
8 h
10 h
KDa
36
IκBα
36
GAPDH
B
***
**
1.2
1
0.8
0.6
0.4
0.2
0
Fold of IκBα expression
Unstimulated
0.5 h
1 h
2 h
4 h
6 h
8 h
10 h
+ (LPS+IFN-γ)
C
100
80
60
40
20
0
Amount of NO production (μM)
Unstimulated-24 h
0.5 h
1 h
2 h
4 h
6 h
8 h
10 h
12 h
24 h
+ (LPS+IFN-γ)

Figure S4

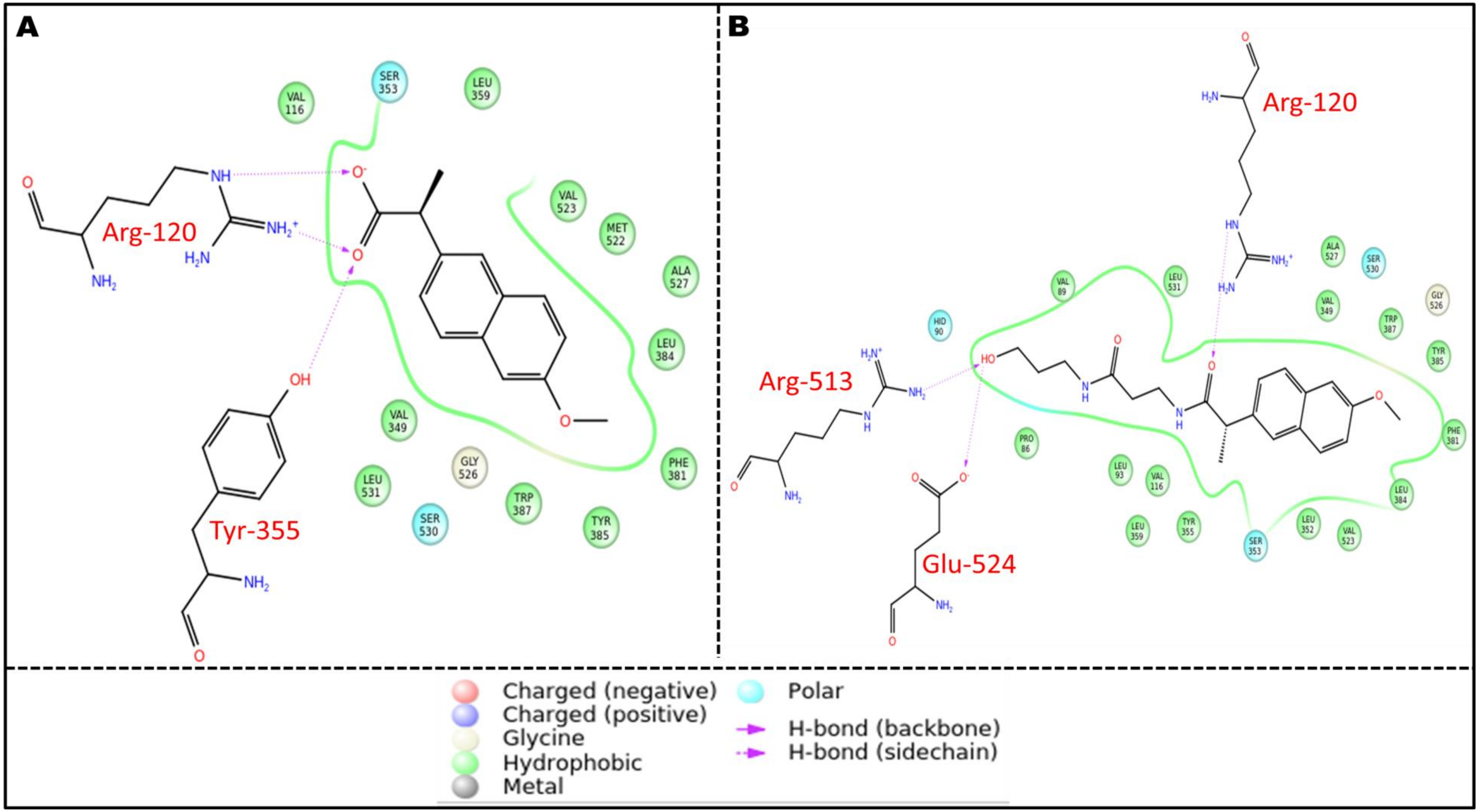
A
B
Arg-120
Tyr-355
Arg-513
Glu-524
Arg-120
Charged (negative)
Charged (positive)
Glycine
Hydrophobic
Metal
Polar
H-bond (backbone)
H-bond (sidechain)

Figure S5
